# Acoustic parameter combinations underlying mapping of pseudoword sounds to multiple domains of meaning: representational similarity analyses and machine-learning models

**DOI:** 10.1101/2024.09.27.615393

**Authors:** G. Vinodh Kumar, Simon Lacey, Josh Dorsi, Lynne C. Nygaard, K. Sathian

**Affiliations:** Departments of Neurology, Penn State College of Medicine, Hershey, PA 17033, USA; Neuroscience & Experimental Therapeutics, Penn State College of Medicine, Hershey, PA 17033, USA; Department of Psychology, Penn State College of Liberal Arts, University Park, PA 16802, USA; Department of Psychology, Emory University, Atlanta, GA 30322, USA

## Abstract

In spoken language, iconicity, referring to the resemblance between the sound structure of words and their meaning, is often studied using pseudowords. Previously, we showed that representational dissimilarity matrices (RDMs) of the shape ratings of pseudowords correlated significantly with RDMs of acoustic parameters reflecting spectro-temporal variations; the ratings also correlated significantly with voice quality parameters. Here, we examined how perceptual ratings relate to these parameters of pseudowords across eight meaning domains. We largely replicated our previous findings for shape, while observing different patterns for other domains. Using a k-nearest-neighbor (KNN) machine-learning algorithm, we compared 4095 combinations of twelve acoustic parameters (3 spectro-temporal and 9 characterizing vocal quality) to determine the optimal combination associated with iconicity ratings in each domain. We found that iconic mappings were linked to domain-specific combinations of acoustic parameters. One spectro-temporal parameter, the fast Fourier transform, contributed to all domains, indicating the importance of time-varying spectral properties for iconicity judgments. We applied the KNN approach to generate shape ratings for 160 real words. These generated ratings strongly correlated with perceptual ratings of real words, indicating the value of the KNN approach to assess iconic mapping in natural languages. Our findings support the relevance of iconicity to language.

## I. INTRODUCTION

Interest in iconicity in spoken language (also known as sound symbolism), that is, the non-arbitrary link between speech sounds and meaning, has been on the rise in the past two decades, challenging the traditionally held view of the ‘arbitrariness of the sign’ in natural languages^1,2^. Indeed, in addition to onomatopoeia, where the sound of the word imitates the sound it describes, sound-meaning mappings have been shown to extend to several non-auditory sensory domains like shape^3,4^, size^5^, weight^6^, and brightness^7^; as well as to non-sensory, abstract, domains, such as valence^8^, arousal^9^, and social dominance^10^. Moreover, non-arbitrary correspondences between sound and meaning have been found at different developmental stages^11,12^, and in various linguistic, geographical and cultural contexts^13^. While converging lines of evidence establish the prevalence and robustness of iconicity in natural languages, it remains unclear how various acoustic features combine and contribute to sound-meaning associations in different domains of meaning.

A major focus of previous research has been on the contributions of phonetic and articulatory features to iconicity^5,14–17^. For instance, pseudowords containing the phonemes /i/ and /e/ connote smallness^5^, brightness^18^, and sharpness^17^, compared to pseudowords containing the vowels /o/ and /u/ that have been associated with roundedness^19^, heaviness^20^ and social dominance^10^. Similarly, the occurrences of certain consonants versus vowels ^21^, voiced versus unvoiced consonants^19,22^, obstruents versus sonorants^19^, or vowel formants^23^ underlie inherent biases in assigning meaning to certain sounds (e.g., rounded versus pointed and dark versus bright). In a related study, we show that perceptual ratings for a large set of 537 pseudowords exhibit domain-specific associations with specific phonetic features, across eight iconic meaning domains^9^. These associations are important since the sound structure of any language is defined by such features.

Whether and how acoustic properties, of either real words or pseudowords, contribute to iconic mappings has received less attention^23^. Acoustic properties may be important because they relate to the listener’s end of the ‘speech chain’^24^, with the spectro-temporal characteristics of speech mapping to phonetic features. For example, intracranial recordings show that, during listening to natural speech, the superior temporal gyrus shows phonetic selectivity arising from neuronal populations tuned to the specific spectro-temporal profiles of each phonetic feature^25–27^. Accordingly, cortical responses to phonetic features of speech could be decoded from low-level acoustic parameters^28^. Additionally, parameters obtained by acoustic analysis have the advantage of defining speech more fundamentally than characterizations relying on linguistic features, enabling the comparison of results from studies across different languages and cultures^23,29^. More importantly, the multidimensional space provided by different acoustic parameters offers a detailed characterization of speech sounds, compared to the higher-level characterization of phonetic and articulatory spaces^30^. Thus, defining iconicity in terms of acoustic parameters complements analyses of phonetic features alone. Few studies have examined the role of different acoustic features in iconicity, either focusing on sound-to-meaning mapping in one or two domains, or employing only a small set of pseudowords^23,29,31^. Consequently, the relative importance of different acoustic parameters in mediating iconic mapping across diverse meaning domains remains largely unknown.

While identifying the key acoustic drivers of iconicity is important, an equally crucial open question is the extent to which these acoustic parameters function within the lexicons of natural languages^32,33^. Since most studies have employed pseudowords to investigate the importance of different phonetic, articulatory and acoustic parameters in mediating iconic associations, it is uncertain to what extent the findings from such studies may apply to the vocabularies of natural languages. Moreover, research of this kind is characterized by a diversity of methods. One approach has been to analyze whether real words conform to the findings for pseudowords, e.g., whether real words for small and large objects contain the same small-associated (high front vowels like /i/) and large-associated phonemes (low-back vowels like /ɑ/), respectively^34^, that have been found in studies using pseudowords. The results of this approach have been limited, with some studies finding clear systematic relationships between vowels in words and their meaning^23,34^, while others either did not find associations^35,36^ or found cross-linguistic and cultural differences, such as the reversal of vowel–meaning associations. The differences across studies further complicate the application of this approach to real words^37^. Another approach employs tasks such as the best-worst judgement format in which participants, on each trial, are presented with a subset of items (e.g., six items) from a list and are instructed to select the two extreme items on a given dimension (e.g., most rounded and most pointed or softest and hardest) in the presented subset^38^. While this method provides a quantitative score of word-meaning association, it is based on ranking of items in particular lists and is thus of limited generalizability. Bridging this gap requires not only identifying the key acoustic parameters underlying iconic associations, but also determining whether such patterns hold in the more complex and variable context of natural language.

Therefore, the purpose of this study is threefold: (1) To provide evidence for the role of acoustic properties in iconic mapping by replicating our previous work in the shape (rounded/pointed) domain and more critically, extending it to seven additional domains: size (small/big), brightness (bright/dark), hardness (hard/soft), roughness (rough/smooth), weight (light/heavy), arousal (calming/exciting), and valence (good/bad); (2) Of primary theoretical interest, to identify the combination of acoustic parameters that best supports iconic associations across the aforementioned eight meaning domains; and (3) As a proof of concept, to use these insights to generate sound iconicity ratings for a set of real words based solely on their acoustic properties, independent of semantic bias. To these ends, we compared perceptual ratings for 537 pseudowords to their corresponding spectro-temporal and vocal parameters, using representational similarity analyses (RSA) for the spectro-temporal parameters and conventional correlation analysis for the vocal parameters, as in a previous study from our group^29^. Next, we employed a *k*-nearest-neighbor (KNN) machine-learning approach to identify the combinations of acoustic parameters that best predicted the perceptual ratings obtained on eight meaning domains. Our approach is based on the principle that, if the acoustic properties of pseudowords vary systematically in accordance to their degree of sound-symbolic association in a given meaning domain, e.g., shape, then similarly rated pseudowords should form clusters in a multidimensional acoustic parameter space. We used this premise to iteratively explore all possible multidimensional acoustic parameter spaces based on the parameters we studied, and find the combinations, in each domain, for which similarly rated pseudowords formed the most coherent clusters, i.e., with minimal distances between such pseudowords. RSA was conducted independently of the combinations identified through our machine-learning approach. Taken together, this set of approaches allowed us to examine the relationship of each parameter to iconicity ratings in each meaning domain, and also identify which parameter combinations best account for iconic associations across different meaning domains. Finally, we extended this approach to generate predicted shape ratings of real words and compare them to perceptual ratings, assessing iconicity while attempting to minimize confounds from semantic processing.

## **II.** METHODS

### A. Participants

As detailed in the report of a related study^9^, participants were recruited and compensated through the Prolific online participant pool^39^ (https://prolific.com). Eligibility criteria included being between 18 and 35 years old and a speaker of American English as their first language, without a history of language or hearing disorders. The sample for seven meaning domains consisted of 389 participants (171 male, 208 female, 6 non-binary, 2 agender, 1 gender-fluid, and 1 who chose not to disclose their gender), with a mean age of 27 years and 1 month (SD = 8 months). The rating tasks were administered via the Gorilla platform^40^ (https://gorilla.sc), where participants also provided informed consent. Participant numbers are reported below for each meaning domain. The procedures received approval from the Emory University Institutional Review Board. For the roughness domain, iconicity rating tasks were administered online to a separate group of 58 participants (26 male, 31 female and 1 non-binary; mean age: 25 years and 9 months, SD = 3 years, 7 months) using the same procedures described above^41^. Participants for these data also provided informed consent, and the study was approved by the Penn State University Institutional Review Board. Additionally, data from 24 monolingual speakers of American English (20 females, mean age 30 years, SD 7 years) recruited for a different experiment^42^ were used in this study. These participants were emailed a link to the experiment on Pavlovia.org after they gave consent over the telephone to participate; this study was approved by the Penn State University Institutional Review Board.

### **B.** Stimuli

We used two sets of audio recordings comprising (1) pseudowords, and (2) real words. As previously reported^29^, the pseudowords were a set of 537 two-syllable CVCV (i.e., consonant/vowel/consonant/vowel) pseudowords created by McCormick et al.^19^, comprising only phonemes and combinations of phonemes that occur in American English^19^. The pseudowords were recorded by a female speaker whose first language was American English, in random order and with neutral intonation, and digitized at a 44.1 kHz sampling rate. Each pseudoword was then down-sampled at 22.05 kHz, which is standard for speech, and amplitude-normalized using PRAAT speech analysis software^43^. The mean duration of the pseudowords was 457 ms (SD ± 62 ms). We also chose 160 real words from Sidhu et al. (2021)^33^, who assessed the meanings of ∼1700 English nouns on a rounded-to-pointed shape dimension. The words were chosen based on their meaning scores such that half came from the two extremes (40 each from the most rounded and the most pointed meaning scores) and the other half from around the median. These words were recorded by a different female speaker of American English. Real-word recordings were amplitude-normalized in Audacity by removing DC offset and setting the peak amplitude to –4.48 dB, with both stereo channels scaled together to maintain their relative levels and spatial balance. Speaker normalization was not performed. The real words had a mean duration of 569 ms (SD ± 117 ms)^42^.

### **C.** Perceptual rating tasks

Participants recruited via Prolific were randomly assigned to rate 537 pseudowords on one of two 7-point Likert type scales representing categorical opposites across eight meaning domains: shape (rounded/pointed, N = 30/30 respectively), size (small/big, N = 32/31), brightness (bright/dark, N = 24/26), hardness (hard/soft, N = 27/27), weight (light/heavy, N = 29/26), arousal (calming/exciting, N = 30/28), and valence (good/bad, N = 26/23)^9^. For the roughness domain (rough/smooth, N = 29/29), participants were recruited following the same procedure as part of another study^41^. We converted these categorically opposing scales in each meaning domain to a single scale by subtracting the ratings given to pseudowords on the roundedness, smallness, brightness, hardness, roughness, lightness, excitement and goodness scales from 8. This operation rescaled the order of the ratings on these scales, i.e., ratings 1 to 7 became 7 to 1, concordant with the order on their corresponding opposing scales. This also yielded a unified scale ranging from 1 to 7, consistently anchored from one categorical extreme to the other, e.g., for shape, the rescaled rating 1 denoted ‘most rounded’ and 7 denoted ‘most pointed’. These unified scales were used to rank the pseudowords along a continuum from one categorical extreme to the other within each meaning domain (see Section II.E.1) and then identify the optimal acoustic parameter combinations in our machine-learning algorithm (see Section II.F).

For rating the 160 real words, 24 participants (on Pavlovia) were randomly assigned to rate their sounds using one of two 7-point Likert type scales; either a 1 (very rounded) to 7 (very pointed) scale or the reverse. Thus, 12 participants provided ratings on each scale. These two sets of ratings were then converted into a unified scale, with 1 representing the most rounded and 7 representing the most pointed, following the procedure described above. For both pseudowords and real words, each participant rated all stimuli once, with the order of presentation randomized. Participants were compensated for their time, with their participation in the pseudoword ratings study lasting approximately 2 hours, and in the real word rating study about 1 hour. To ensure data quality, both tasks incorporated attention checks consisting of responding to tasks unrelated to the primary rating task, e.g. what is the color of a carrot?

### **D.** Acoustic parameters

Following our prior study^29^, we selected three spectro-temporal acoustic parameters: the fast Fourier transform (FFT), spectral tilt, and speech envelope, measuring the distributions of frequency and amplitude over time^44,45^; and seven parameters capturing the degree and regularity of voicing. These included 3 parameters indexing the proportion and characteristics of voiced segments^43^: the fraction of unvoiced frames, mean autocorrelation (MAC), and pulse number; 3 parameters capturing the regularity of voicing or voice quality (hereafter: voice parameters) during the production of each pseudoword: jitter, shimmer, and harmonics-to-noise ratio (HNR)^46^; and the standard deviation of the fundamental frequency of the speech sound (F0_SD_)^47^. We also included 2 additional parameters: the mean F0, and item duration. Each parameter is described below.

### 1. Spectro-temporal parameters

a. *Fast Fourier transform (FFT)*

The FFT extracts the frequency components of the speech signal and captures how their energy changes over time, thereby representing the power spectrum throughout the duration of the spoken pseudowords. The FFT is visualized through the spectrogram, which displays the distribution of power across frequencies over time. For instance, obstruents are typically reflected in sudden changes in power across different frequencies, whereas sonorants show more gradual variations in power across frequencies (see spectrograms in **Supplementary** Figure 9b).

*Spectral tilt*

Spectral tilt captures relative differences in power across frequencies, providing an estimate of the overall slope of the power spectrum based on measurements taken across the entire utterance. When lower frequencies contain more power than higher ones, the power spectrum shows a steep downward slope from low to high frequencies. This slope becomes flatter when more power is spread across both lower and higher frequencies.

*Speech envelope*

The speech envelope captures fluctuations in amplitude over time, closely aligning with phonemic features and syllabic transitions^45^. When these transitions are abrupt, such as with stops, affricates, or fricatives, the envelope becomes irregular and discontinuous. In contrast, smoother and more gradual transitions, as seen with sonorants, result in a more continuous and uniform envelope.

### 2. Vocal parameters

a. *Mean autocorrelation (MAC)*

This is a measure of the periodicity of a signal^29,43^. Periodicity is expected to be high for a long vowel such as /u_ː_/ or a consonant like /m_ː_/ where successive periods should be very similar, i.e., highly correlated. Higher autocorrelation values reflect smoother, more regular voicing and a greater presence of voiced segments, while lower values indicate a more irregular pattern and fewer voiced or periodic segments.

*Pulse number*

This refers to the number of glottal pulses, meaning the opening and closing of the vocal folds, that occur during the production of vowels or voiced consonants, measured across the entire utterance^29,43^. A lower pulse number suggests a more uneven voice pattern or fewer voiced segments, whereas a higher pulse number suggests a smoother voice pattern or more voiced segments.

*Fraction of unvoiced frames*

The fraction of unvoiced frames indicates the proportion of time windows in which the vocal folds are not active, expressed as a percentage^29,43^ . The fraction of unvoiced frames is influenced by the phonemic composition of a pseudoword — higher values are observed in pseudowords containing unvoiced features, such as voiceless stop consonants, while lower values are typical for those with voiced, periodic elements like vowels.

*Shimmer*

Shimmer measures the variability in peak-to-peak amplitude of the glottal waveform^48^ . It serves as an indicator of vocal stability, with low shimmer corresponding to smooth, stable speech and high shimmer indicating irregular amplitude fluctuations, often perceived as hoarseness in the voice.

*Jitter*

Jitter refers to the variation in frequency between successive vocal cycles and serves as an indicator of voice quality by capturing irregularities in vocal fold vibration^46^. High jitter values are associated with a rough or “breaking” voice quality. Jitter is typically assessed using sustained vowel sounds, where minimal frequency variation is expected. In speech production, elevated jitter indicates greater vocal instability and contributes to the perception of vocal roughness.

*Mean harmonics-to-noise ratio (HNR)*:

This ratio reflects the relationship between the periodic (or harmonic), and aperiodic (or noise), components of the speech signal. The mean HNR provides an estimate of the sound’s overall periodicity, expressed in decibels (dB)^46^. The noise component results from turbulent airflow at the glottis when the vocal folds fail to close completely^49^. As noise increases, and therefore, mean HNR decreases, the voice becomes increasingly hoarse and the speech pattern becomes progressively more uneven^49^.

*F0 & F0_SD_*

F0 is the average fundamental frequency of the speech signal, while F0_SD_ captures the variability in F0 across the utterance^29,43^. F0_SD_ reflects the degree of change in intonation: a low F0_SD_ corresponds to a flat, monotone voice, whereas a high F0_SD_ indicates a more dynamic or expressive vocal tone^47^.

*Duration*

The duration of the speech signal, measured in milliseconds (ms), is influenced by several factors, including the distinction between long and short vowels^50,51^, the surrounding phonetic context^50^, and the speaker’s rate of speech^52^. Although this is not, strictly, a voice quality parameter, it is grouped here with the vocal parameters for convenience, as they are all measured with a single value per item, unlike the spectro-temporal parameters (see below).

### **E.** Analysis of acoustic parameters

**1.** *Spectro-temporal parameter analysis*

RSA, originally developed for analyzing functional magnetic resonance imaging (fMRI) data^53^, was applied in our previous study^29^ to compare perceptual ratings of pseudowords on the rounded-pointed continuum of shape to their spectro-temporal acoustic features. In the present study, we employed a similar RSA approach across all eight domains.

For each domain, we constructed a 537 × 537 representational dissimilarity matrix (RDM) based on participants’ perceptual ratings of the pseudowords. Using the unified scale (see section II.C), pseudowords were ranked by their mean rating along the domain-specific dimension **(Figure 1**, Step 1): from most rounded to most pointed (shape), smallest to biggest (size), hardest to softest (hardness), smoothest to roughest, lightest to heaviest (weight), brightest to darkest (brightness), most calming to most exciting (arousal), and best to worst (valence). Based on these ordered values, we then created reference RDMs for each domain (**Figure 1**, Step 2). This involved calculating the first-order Pearson correlation between the vectorized perceptual ratings of each pair of pseudowords across participants using the original (unrecoded) data, since RDMs capture dissimilarity irrespective of the scale used by any individual. The dissimilarity between each pair was defined as 1 minus the correlation coefficient (1 – r), and this value was entered into each cell of the RDM. Note that, while correlations range from +1 to -1, the corresponding dissimilarity values range from 0 to 2.

**Figure 1:**
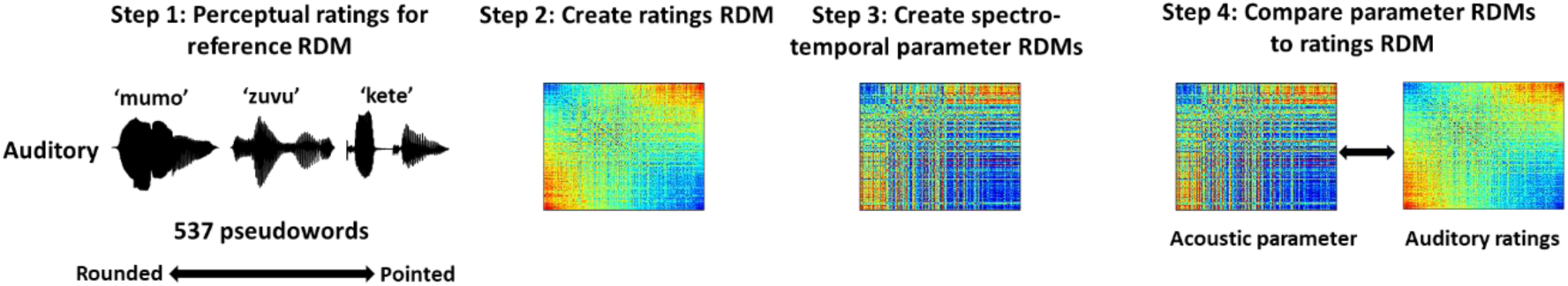
**Schematic illustration of the analysis pipeline**. Step 1: auditory pseudowords were rated along different dimensions, e.g., shape (rounded/pointed), brightness (bright/dark). Step 2: these perceptual ratings were used to create reference representational dissimilarity matrices (RDMs). Step 3: RDMs for each of the spectro-temporal parameters were created for each domain. Step 4: in each domain, the spectro-temporal parameter RDMs were compared to the ratings RDM by way of a second-order correlation.

We also created corresponding acoustic RDMs (**Figure 1**, Step 3) by vectorizing the spectro-temporal measurements for each pseudoword, computing pairwise correlations between items, and converting these to dissimilarity values using the 1 – r metric. These acoustic RDMs were computed separately for the speech envelope, spectral tilt, and FFT. Spectro-temporal parameters and their associated RDMs were computed in MATLAB (version 2021a; The MathWorks, Natick, MA), following the procedures outlined earlier^29^ . To ensure consistency across items, we normalized the duration of all pseudowords to the mean duration of 457 ms using MATLAB’s resampling function. At a down-sampled rate of 22.05 kHz (see section II.B), each pseudoword was represented by a common vector length of 10,077 data points (22,050 × 0.457). This normalization introduced only minimal noise, proportional to the relative variability in duration (13.5%, calculated as SD/mean = 62/457 ms). Standardizing the length allowed for equal dimensionality across pseudowords in the RDM computations. It is worth noting that the spectro-temporal parameters are not all independent, e.g., both spectral tilt and FFT capture aspects of the distribution of frequency content, and that each parameter comprises multiple values per pseudoword. The exact measurements extracted from these vectors varied across parameters, for example differing window lengths and overlaps (details provided in the Supplementary Material). As a result, the number of data points underlying the dissimilarity calculations differed by parameter. For instance, the speech envelope and FFT had 10077 and 5489 data points respectively, and pairwise dissimilarities were computed using Pearson correlations. In contrast, spectral tilt was represented by only 8 data points, so dissimilarities were computed using Spearman (non-parametric) correlations.

To assess the relationship between the perceptual and acoustic representational structures, we compared each domain-specific ratings RDM with each of the corresponding spectro-temporal RDMs (Figure 1, Step 4). For each comparison, we vectorized the lower triangular portion (excluding the diagonal) of both RDMs and computed a second-order Spearman correlation between the two vectors, yielding the observed correlation coefficient (ρ). Statistical significance was assessed using a non-parametric permutation test: the elements of the vectorized ratings RDM were randomly permuted, and the Spearman correlation with the fixed acoustic RDM was recomputed 10,000 times to generate a null distribution of correlation values. The p-value was then calculated using the equation below, providing an estimate of the probability that the observed relationship arose by chance. Significant correlations at this level indicate which spectro-temporal features are associated with perceptual judgments of the pseudowords.

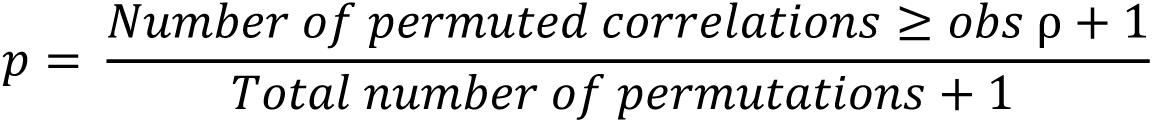

**2.** *Voice parameter analysis*

As reported previously^29^, the voice parameters were measured in PRAAT^43^, using the original pseudoword sound files, without applying the resampling procedure described earlier. Similar to the spectro-temporal parameters, the voice parameters are not necessarily independent of one another. For example, the fraction of unvoiced frames and pulse number both measure the proportion of an utterance that is voiced. However, unlike the spectro-temporal parameters, the voice parameters are represented by a single value for each pseudoword. For calculating jitter and shimmer, we used the local measures, which represent the mean absolute difference in frequency and amplitude, respectively, between consecutive periods of the speech waveform. These differences were then divided by the overall mean difference across all periods and expressed as a percentage^43^. Note that the average duration of the vowel segments in our study was 119 ms for vowel 1 and 238 ms for vowel 2, well above the 50 ms threshold typically used to ensure reliable measurements^54,55^. Moreover, our acoustic analyses were conducted on the entire pseudoword, meaning that the stretches of periodicity used by PRAAT to compute measures such as jitter and shimmer consistently exceeded 50 ms for all items (and indeed for each syllable within them). Each voice parameter, represented by a single value per pseudoword, was directly compared to the recoded perceptual ratings using standard correlational analyses. Since we examined correlations between ratings and nine voice parameters, the significance of each correlation was assessed after Bonferroni correction (corrected α = 0.0056).

### **F.** *k*-nearest neighbor models

A *k*-nearest neighbor (KNN) machine-learning algorithm was employed to find the set of acoustic parameters that correlated best with the unified ratings (see section II.C) for the 537 pseudowords in each of the eight meaning domains studied. The algorithm (**Figure 2**) comprised five steps as described below.

1. Step 1: Data preparation

**Figure 2:**
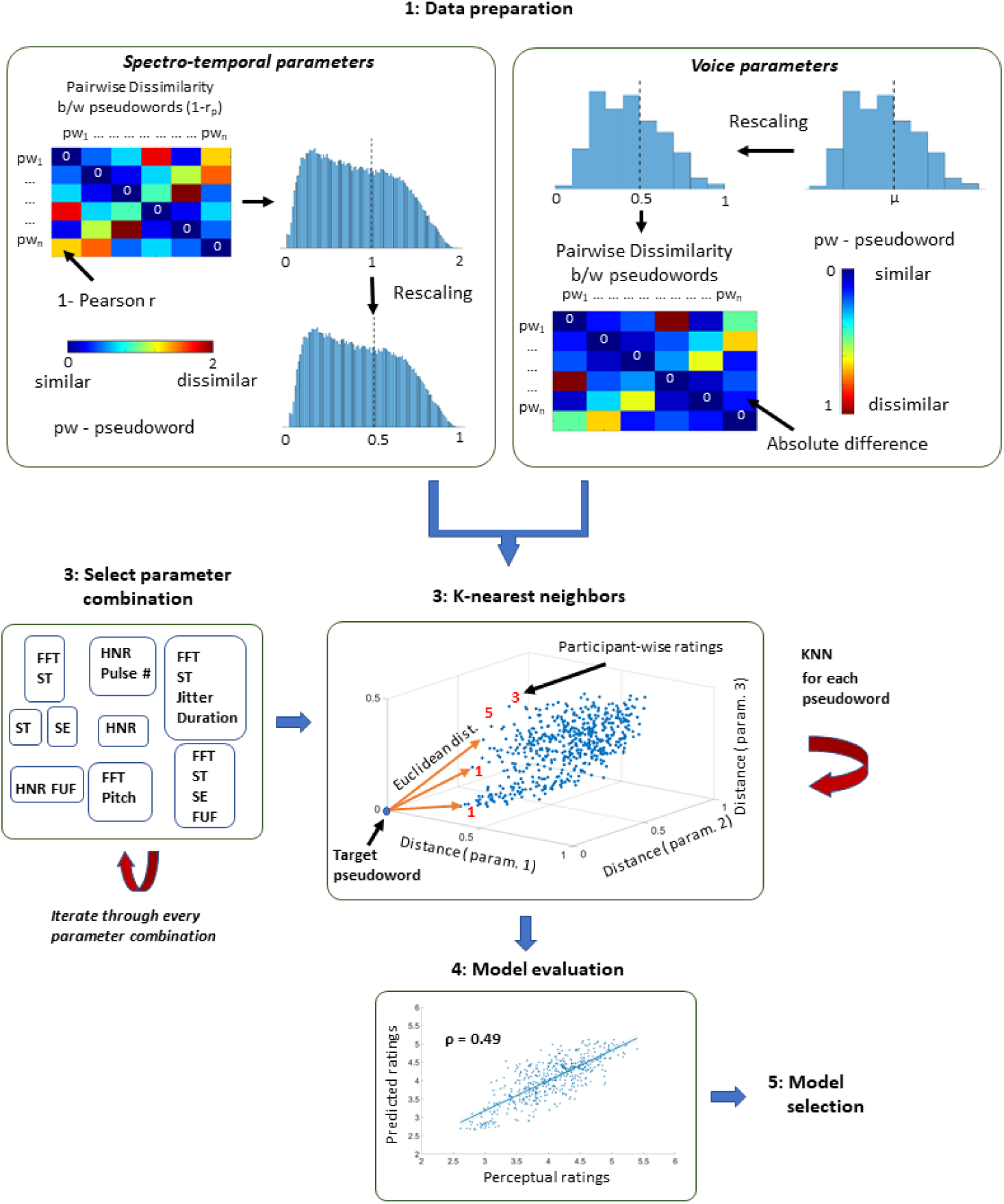
Analysis flowchart for generating pseudoword ratings based on acoustic parameters. Step 1: The pairwise distance between pseudowords was computed based on their differences in spectro-temporal and voice parameters, as detailed in the text. **Step 2.** The number of parameters entering the algorithm was varied from 1-12 and all 4095 combinations were iteratively tested. **Step 3.** For each combination, Euclidean distance was iteratively computed between each pseudoword and the remaining 536 pseudowords in a n-parameter space. The predicted ratings of each pseudoword were defined by the modal perceptual rating of its closest 23 neighbors (k = √537 ≈ 23). **Step 4.** Spearman correlations (ρ) between the perceptual and model-based ratings on a participant-by-participant basis were used to quantify the performance of the model. **Step 5.** The parameter combination that yielded the highest ρ was chosen as the best model for the corresponding meaning domain. ratings of these neighbors vary across participants. Consequently, each real word’s rating was predicted separately for each participant. These participant-specific predicted ratings were then averaged across all participants, and the mean predicted ratings were correlated with the mean perceptual ratings for each item using Pearson’s correlations. We acknowledge that participants’ perceptual ratings for real words may not be entirely free from semantic influences, even though the participants were instructed to respond based solely on the sounds of the real words. However, the perceptual ratings of the real words were used only as a means to validate our findings.

Each parameter extracted for a particular pseudoword served as a separate dimension on which pairwise distances were computed between pseudowords. Pairwise distances between pseudowords were computed for these parameters using the methods described in section II.E.1. The dissimilarity values derived from the spectro-temporal parameters were then linearly rescaled to range between 0 and 1. For the nine voice parameters, each pseudoword was represented by a single value. Pairwise distances between pseudowords for these parameters were computed by linearly rescaling the parameter values to lie between 0 and 1 and then taking the absolute difference between the corresponding values of each pseudoword pair. The rescalings were performed to ensure that the pairwise distances between pseudowords based on the spectro-temporal and voice parameters were on a uniform scale.

2. Step 2: Selection of parameter sets for testing

Since the primary goal of the analyses was to identify the set of acoustic parameters that best correlated with the ratings of the pseudowords, we varied the number of parameters from 1-12 and iteratively performed the subsequent steps with all 4095 possible parameter combinations.

3. Step 3: k-nearest neighbors

The KNN model rating for every pseudoword was iteratively generated in each of the multidimensional parameter spaces defined by a particular combination of acoustic parameters. In each of these spaces, the computed rating for a given pseudoword depends on the perceptual ratings of its nearest neighboring pseudowords. The distances between pseudowords were determined by Euclidean distances while neighborhood size was taken as the square root of the total number of pseudowords: k = √537 ≈ 23 ^Ref^ ^56^. In each parameter space, the rating for a given pseudoword was computed based on the modal ratings of its 23 nearest neighbors (for instance, if a given pseudoword’s neighbors were most frequently rated as 3, then the given pseudoword was assigned the predicted rating 3). The procedure was repeated iteratively for every pseudoword. It is important to note that, in each parameter space, the 23 nearest neighbors for a given pseudoword are invariant because the Euclidean distances between pseudowords are based on their acoustic features, whereas the ratings of these neighbors vary on a participant-by-participant basis. Thus, the rating for each pseudoword was computed on a participant-wise basis in all 4095 parameter spaces (N for each domain: shape, 60; size, 63; brightness, 50; hardness, 54; roughness, 58; weight, 55; arousal, 58; valence, 49).

### Steps 4 and 5: Model evaluation and selection

In each meaning domain, participant-wise computed ratings for 537 pseudowords were concatenated into a vector for each combination of acoustic parameters and correlated with the corresponding vectorized perceptual ratings. Because of the ordinal nature and non-normal distribution of the ratings, Spearman’s correlation (ρ) was used. The combination of parameters yielding the highest ρ value for a given meaning domain was chosen as the optimal combination for computing model-based pseudoword ratings in that domain.

### **G.** Shape ratings of real words

As a proof of concept, we sought to apply our approach to predict the shape ratings of real words while attempting to minimize the impact of semantic knowledge. Shape ratings for real words were based on the perceptual ratings of pseudowords in the acoustic neighborhood of each real word^42^. Using the KNN procedure outlined in Section II.F, we determined the most relevant acoustic parameters for classifying the shape ratings of pseudowords. These parameters defined the multidimensional acoustic space within which the ratings for real words were estimated. Each real word was iteratively projected into this space, and its 23 nearest pseudoword neighbors (k = √538 ≈ 23) were determined using Euclidean distances from all 537 pseudowords. The modal shape rating of these neighbors was then assigned to the real word, as it reflects the sound-symbolic judgment derived from the perceptual ratings of pseudowords occupying the same acoustic neighborhood. As outlined in Section II.F.3, the set of 23 nearest pseudoword neighbors for each real word remains invariant because the Euclidean distances to the pseudowords are determined solely by their acoustic features (i.e., the optimal parameter combination for shape). However, the

In addition to the aforementioned analyses, we performed three supplementary procedures to examine perceptual variability across participants and assess the robustness of our findings. We first quantified participant-level variability in perceptual ratings by calculating confidence intervals for each pseudoword. Next, we explored the effect of varying *k* in the KNN algorithm across semantic domains to assess the sensitivity of model performance to parameter choice. Finally, we conducted subsampling analyses to evaluate the stability of feature selection across participant subsets. Detailed descriptions of the methods and results for these analyses are provided in the Supplementary Materials.

## **III.** RESULTS

### **A.** Acoustic parameters mediating iconic associations for each meaning domain

For each domain, the following sections present: RSA for individual spectro-temporal parameters, conventional correlations between each voice parameter and the perceptual ratings, and the optimal parameter combination identified by the KNN model.

1. Shape
a. *RSA for spectro-temporal parameters*

Replicating our previous findings^29^, RSA showed that the RDMs for all three spectro-temporal parameters were significantly positively correlated with the RDM for ratings of the pseudowords from rounded to pointed (**Figure 3B-D**). The strongest correlation was for spectral tilt (r = .41, p < .0001), which was steeper for pseudowords rated as more rounded where power was concentrated in low-frequency bands, but flatter for pseudowords rated as pointed in which power migrated to the higher-frequency bands (see representative highly rated rounded and pointed pseudowords in **Supplementary** Figure 9a). This pattern likely reflects underlying phonetic contrasts in sonorant versus unvoiced obstruents, as well as the influence of place of articulation, with bilabial and labiodental consonants contributing to lower-frequency spectra and alveolar or velar consonants contributing to higher-frequency components^9^. The next strongest relationship was for the FFT (r = .28, p < .0001), which indicates that FFT patterns changed more gradually over the duration of the pseudoword for rounded-rated pseudowords and more abruptly for pointed-rated ones, further distinguishing, e.g. the smoothness of sonorants from the sharp onsets of obstruents (**Supplementary** Figure 9b). The speech envelope (r = .18, p < .001) showed a more modest effect. It was smoother and more continuous for pseudowords rated as strongly rounded, reflecting sustained vocal energy and reduced temporal variability, whereas those rated as strongly pointed had a more discontinuous and uneven envelope (**Supplementary** Figure 9c).

*Conventional correlations for voice parameters*

**Figure 3:**
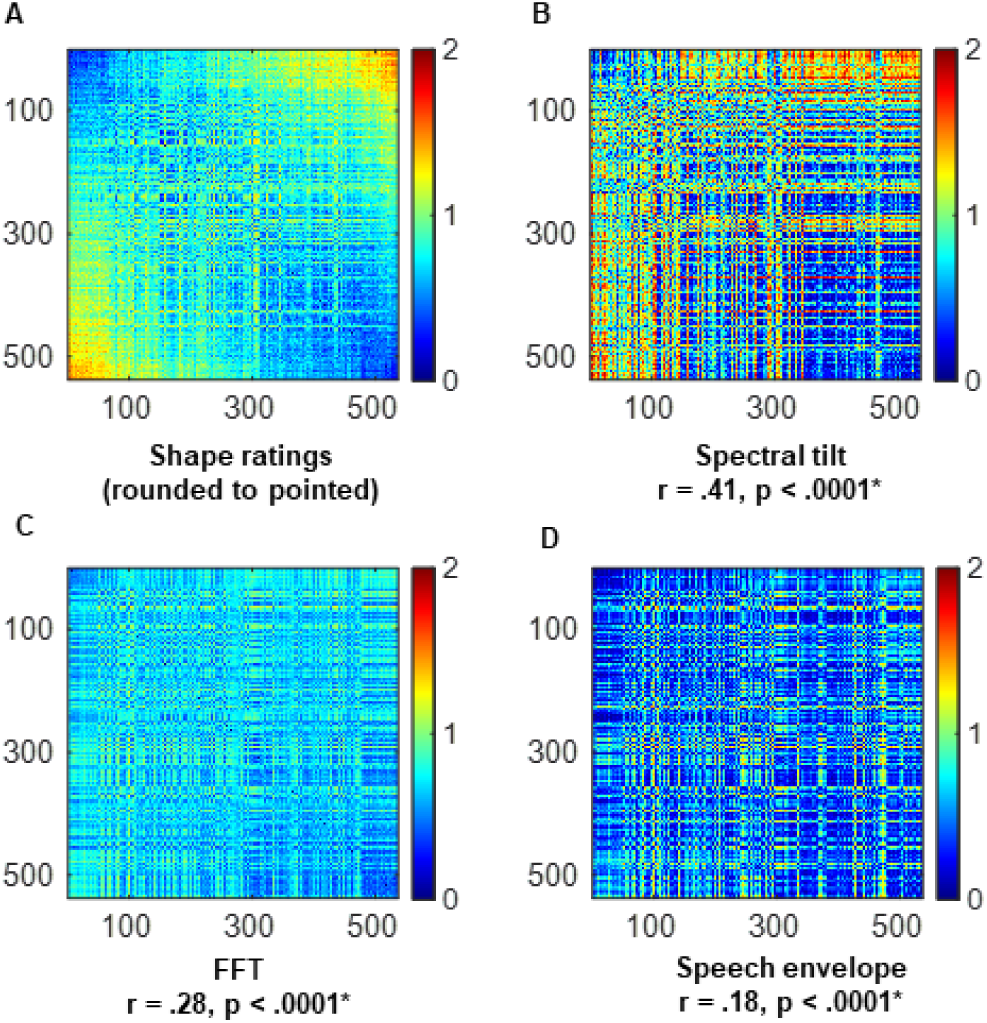
Representational similarity analyses relating auditory shape ratings and spectro-temporal parameters. **A**: Ratings RDM. **B–D**: RDMs for the three spectro-temporal parameters. Pseudowords are ordered, left to right, from most rounded to most pointed. Spectro-temporal parameter RDMs are ordered (top right, bottom left, bottom right) from the one with the strongest to the one with the weakest relationship. Color bar shows pairwise dissimilarity (1-r), where 0 = zero dissimilarity (items are identical) and 2 = maximum dissimilarity (items are completely different). ρ = Spearman correlation coefficient for second-order correlation between the parameter RDM and ratings RDM; significance was derived using a permutation-based Mantel test (10,000 iterations).

For the voice parameters (**Table 1** and **Supplementary** Figure 10a-i), the mean HNR (r = -.58, p<0.001), pulse number (r = -.48, p<0.001), and MAC (r = -.42, p<0.001) were significantly negatively correlated with ratings of the pseudowords, reflecting the presence of more sonorants in items rated as rounded compared to those rated as pointed. The fraction of unvoiced frames was significantly positively correlated with ratings (r = .50, p<0.001), reflecting the greater presence of unvoiced segments in items rated as more pointed. Shimmer (r = .39, p<0.001) and jitter (r = .30, p<0.001) were significantly positively correlated with ratings, reflecting increasing variability in voice quality as ratings of the pseudowords transitioned from rounded to pointed. Thus far, the voice parameter results replicated our earlier study^29^ but, in the present study, and in contrast to our earlier study, F0_SD_ was correlated significantly and positively with ratings (r = .17, p<0.001), increasing as ratings moved from rounded to pointed. This indicates greater variability in the fundamental frequency for pointed, compared to rounded, pseudowords. Of the two newly examined parameters, duration (r = -.13, p=0.002) was inversely related to pointedness, aligning with underlying phonetic contrasts in sonorant versus obstruent content for rounded and pointed pseudowords, while F0 (r = .03, p = 0.5) was not significantly correlated with ratings. Together, these findings provide a robust replication of our prior results and offer some new insights into the acoustic dimensions of pseudoword sounds that underlie their iconic associations in the rounded-pointed domain. Importantly, they highlight how increasing acoustic variability, temporal irregularity, and spectral shift toward higher frequencies all contribute to the perceptual transition from rounded to pointed.

*Optimal combination*

**Table 1:**
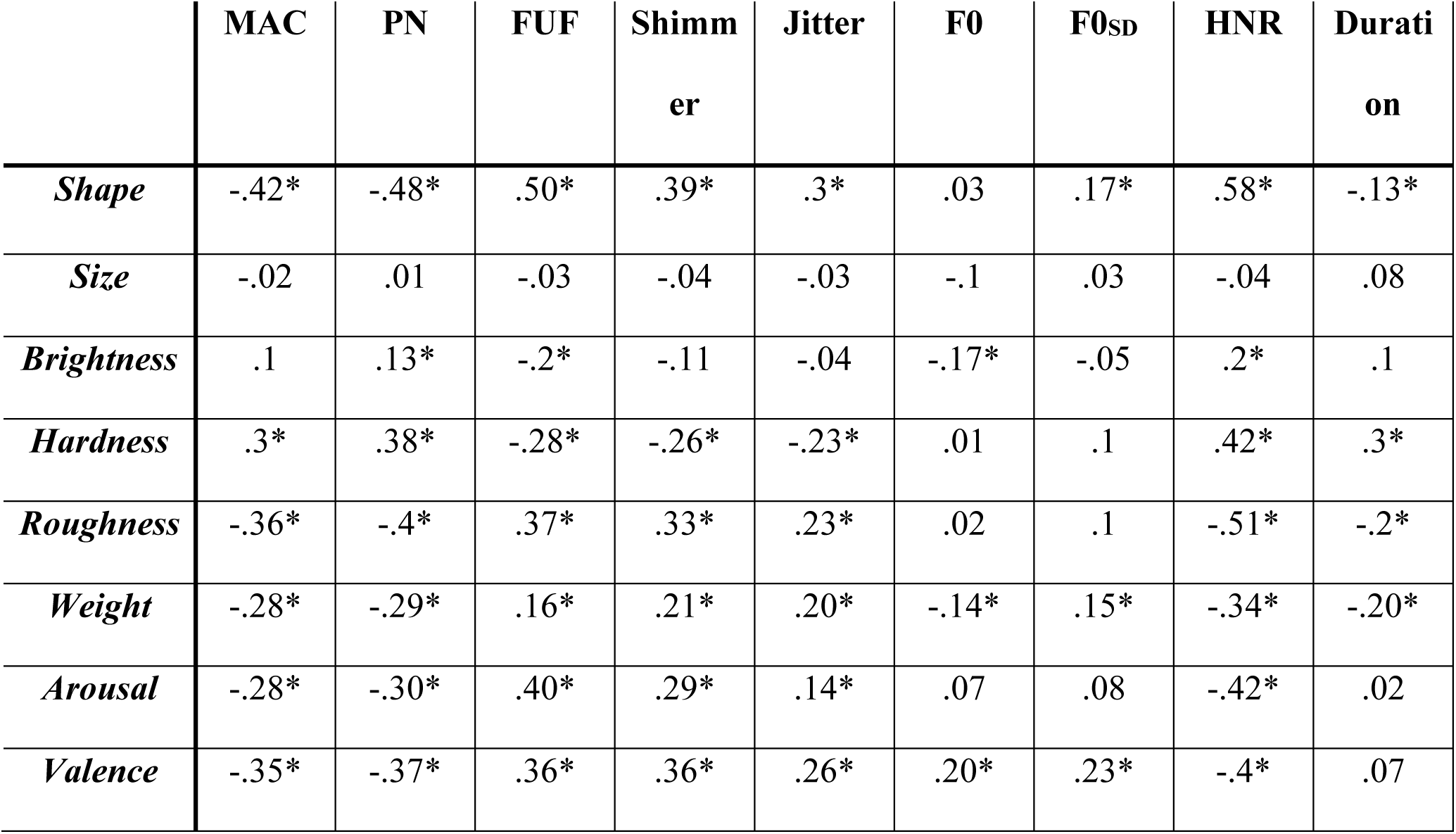
Pearson correlations between auditory perceptual ratings across different meaning domains and nine voice-quality parameters. Pearson; *correlation passes the Bonferroni-corrected α of 0.0056 for nine tests; df = 535 for all nine voice parameters. PN: Pulse number, FUF: Fraction of unvoiced frames, HNR: Harmonics-to-noise ratio, MAC: Mean autocorrelation

|  | MAC | PN | FUF | Shimmer | Jitter | F0 | F0<sub>SD</sub> | HNR | Duration |
| --- | --- | --- | --- | --- | --- | --- | --- | --- | --- |
| *Shape* | -.42* | -.48* | .50* | .39* | .3* | .03 | .17* | .58* | -.13* |
| *Size* | -.02 | .01 | -.03 | -.04 | -.03 | -.1 | .03 | -.04 | .08 |
| *Brightness* | .1 | .13* | -.2* | -.11 | -.04 | -.17* | -.05 | .2* | .1 |
| *Hardness* | .3* | .38* | -.28* | -.26* | -.23* | .01 | .1 | .42* | .3* |
| *Roughness* | -.36* | -.4* | .37* | .33* | .23* | .02 | .1 | -.51* | -.2* |
| *Weight* | -.28* | -.29* | .16* | .21* | .20* | -.14* | .15* | -.34* | -.20* |
| *Arousal* | -.28* | -.30* | .40* | .29* | .14* | .07 | .08 | -.42* | .02 |
| *Valence* | -.35* | -.37* | .36* | .36* | .26* | .20* | .23* | -.4* | .07 |

While 11 of the 12 parameters examined were found to be significantly related to shape ratings in the preceding analyses (all except F0), the KNN algorithm identified an optimal model comprising only five of these parameters, presumably as a consequence of the different approach. The optimal model included two spectro-temporal parameters: the FFT and spectral tilt, and three vocal parameters: the fraction of unvoiced frames, shimmer, and duration. This combination yielded ratings that showed the highest correlation (ρ = 0.59), among all the combinations tested, with participants’ (N = 60) perceptual ratings of the 537 pseudowords in the shape domain (**Table 2**).

**Table 2:** Summary of KNN results FFT: Fast Fourier transform, ST: Spectral tilt, SE: Speech envelope, PN: Pulse number, FUF: Fraction of unvoiced frames, HNR: Harmonics-to-noise ratio, MAC: Mean autocorrelation

| **Domain** | **N** | **Max. $\rho$** | **Parameter combination in the optimal model** |
| --- | --- | --- | --- |
| *Shape* | 60 | 0.59 | FFT, ST, FUF, shimmer and duration |
| *Size* | 63 | 0.45 | FFT, ST, SE, FUF and duration |
| *Brightness* | 50 | 0.52 | FFT, ST, SE, PN, FUF, jitter, MAC, F0, F0<sub>SD</sub> and duration |
| *Hardness* | 54 | 0.54 | FFT, ST, FUF, shimmer and duration |
| *Roughness* | 58 | 0.57 | FFT, SE |
| *Weight* | 55 | 0.45 | FFT, ST, SE, FUF, HNR and duration |
| *Arousal* | 58 | 0.51 | FFT, ST, SE, FUF and HNR |
| *Valence* | 49 | 0.66 | FFT and SE |

Existing studies show that formant frequencies are higher for more pointed words^23^ and for vocalizations produced in response to more pointed shapes^57^, indicating the significance of formant frequencies in mediating sound-shape relationships. Each formant is associated with a specific frequency range and is characterized by a concentration of spectral power around those frequencies^58^. Formant structure of the pseudowords reflect the resonant frequencies shaped by vocal-tract configuration and directly determine how acoustic energy is distributed across frequencies. Variations in formant frequencies modulate the FFT spectrum by shifting where energy peaks occur and influence spectral tilt by altering how rapidly energy declines from low to high frequencies, e.g. rounded/pointed words have power concentrated at lower/higher frequencies resulting in a steeper/flatter spectral tilt, whereas the FFT reflects the power spectrum across the duration of the spoken pseudowords (**Supplementary** Figure 9a), capturing the instantaneous spectral profile as the pseudoword unfolds in time.

The duration of the speech signal carries information related to the component vowels^50,59^ that influence sound-shape judgments, e.g., rounded (e.g., /o/) and unrounded vowels (e.g., /ɪ/) that influence roundedness and pointedness, respectively, have distinctive temporal profiles. Additionally, the type of vowels that precede or follow a consonant can influence the duration of the coarticulated phoneme and in turn, the syllable, thereby affecting the overall duration of the pseudoword. Importantly, these duration effects are likely not limited to English; duration is a cross-linguistic prosodic cue^60^. For example, final lengthening, where syllables or segments at the ends of words and phrases are universally produced with longer duration^61^, provides listeners with a robust cue for speech segmentation and perception^60,62^. However, the pseudowords in our set were produced with neutral and consistent prosody. Therefore, the observed duration differences are more likely tied to intrinsic vowel–consonant dynamics. Robust sound-shape associations have been shown between sonorants (e.g., /l/, /m/, /n/) and round shapes, and voiceless stops (/p/, /t/, /k/) and pointed shapes^9^, supporting the importance of the fraction of unvoiced frames characterizing features related to the degree of voicing in speech. The presence of shimmer in the model is consistent with the importance of the amplitude profile, as shimmer indexes the peak-to-peak variation in the amplitude of the periodic speech signal, implying that momentary amplitude variations are important in mediating sound-iconic judgements of shape.

2. Size
a. *RSA for spectro-temporal parameters*

RSA revealed weak but significant correlations between the RDM for size ratings and those for spectral tilt (r = .12, p < .0001), speech envelope (r = .11, p < .0001), and FFT (r = .08, p < .0001) (Figure 4B–D). Spectral tilt was broadly similar for pseudowords rated as small or big, with initially steep but flattening slopes. However, big-rated pseudowords retained relatively higher broadband energy (**Supplementary** Figure 11a). Both big-and small-rated pseudowords showed discontinuous speech envelopes, though big-rated pseudowords had slightly greater amplitude in the second syllable (**Supplementary** Figure 11b). Spectrograms revealed comparable energy changes overall, with somewhat greater high-frequency energy for small-rated pseudowords (**Supplementary** Figure 11c). The weak spectral tilt correlation suggests that both small- and big-rated pseudowords comprise a mix of low- and high-frequency energy due to voiced/unvoiced consonants and their places of articulation. In small-rated pseudowords, unvoiced consonants at bilabial/labiodental places balance out to produce mid-range spectral energy, while big-rated pseudowords combine high-frequency fricatives with low-frequency post-alveolar/velar consonants and back rounded vowels to generate a comparable spectral profile^9^. The shared discontinuous envelopes likely reflect stops, whereas the slightly greater second-syllable amplitude in big-rated pseudowords may arise from voicing differences influencing intensity and periodicity, providing subtle cues that may contribute to iconicity judgments in the size domain.

*Conventional correlations for voice parameters*

**Figure 4:**
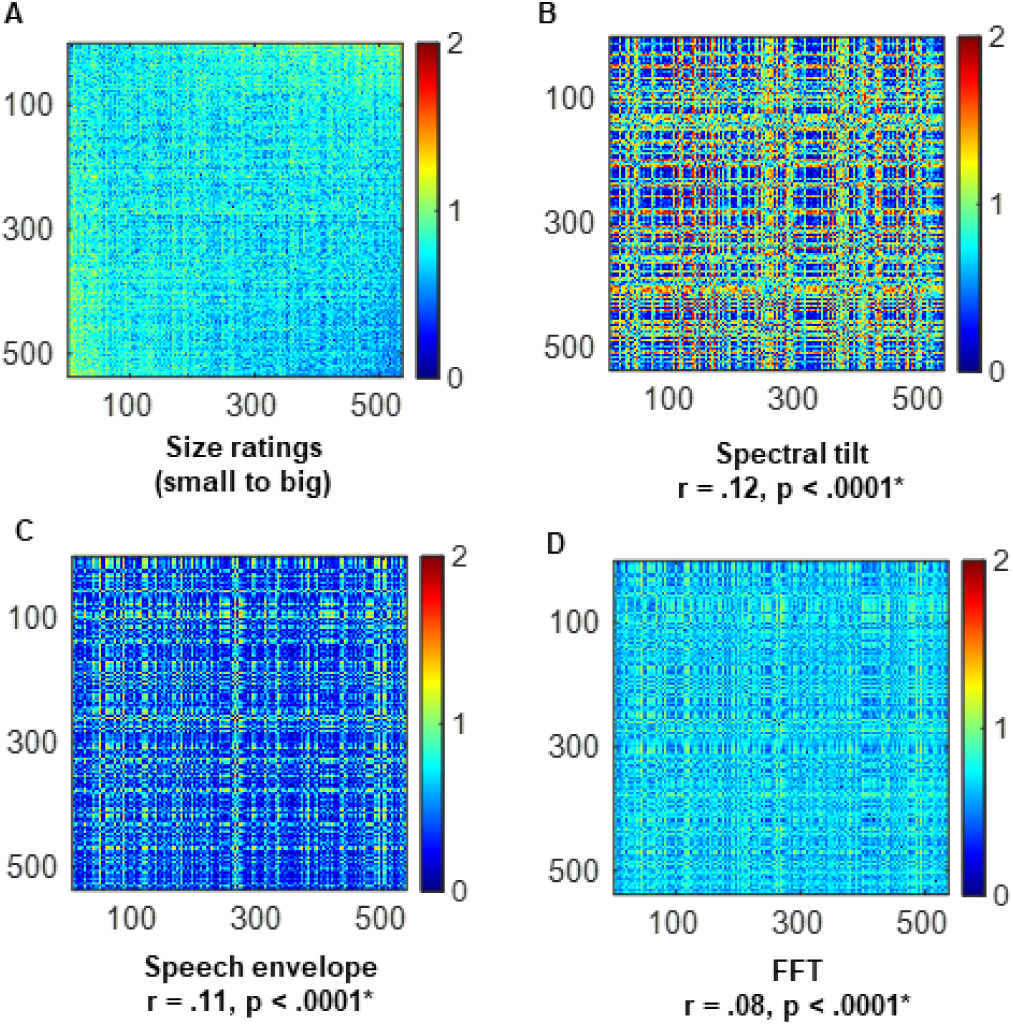
Representational similarity analyses and conventional correlations relating auditory size ratings and acoustic parameters. **A**: Ratings RDM. **B–D**: RDMs for the three spectro-temporal parameters. Pseudowords are ordered, left to right, from smallest to largest. All other details as for Fig. 3

There were no significant correlations between voice parameters and size ratings that survived correction (**Table 1** and **Supplementary** Figure 12a-i). This pattern indicates the inadequacy of conventional correlations to capture the underlying sound-size relationships for this specific pseudoword set. This pattern may reflect characteristics of the particular pseudowords used, since they were selected based on associations of the component phonemes to roundedness and pointedness. A different set of pseudowords incorporating phonemes known to be associated with size iconicity did yield correlations between size ratings and some of the voice parameter^63^. While prior work has linked duration and phonation type to size iconicity^23^, these effects may depend on intentional exaggeration by speakers or specific experimental contexts. In our study, the speaker used natural intonation and voicing, which may have muted such associations. Taken together, these results highlight that while certain acoustic properties may weakly reflect iconic size ratings, they do not provide strong or consistent cues, at least in the present set of pseudowords.

*Optimal combination*

The KNN algorithm showed that perceptual ratings of size of the 537 pseudowords (N = 63) were best correlated (ρ = 0.45) with the ratings predicted by a combination of five parameters. This combination included all three spectro-temporal parameters: the FFT, spectral tilt, and speech envelope, and two voice parameters: the fraction of unvoiced frames and duration (**Table 2**). Although no single vocal parameter showed a significant correlation with size judgments on its own, these parameters may have contributed to the KNN model for size because the model exploits joint patterns across features. Even if each parameter contributes only minimal information individually, the combined interactions can capture more complex relationships that support sound-symbolic judgments.

Evidence suggests that the frequency of the vocalizations produced by animals is related to their size, i.e., animals with larger bodies produce vocalizations with lower frequencies, and vice versa^64–66^. It has been argued that experience of such co-occurring attributes (e.g. body size and frequency profile of the vocalization) in the environment leads to internalization of these probabilities and might underlie iconic associations^17^. Furthermore, small/big ratings were found to be associated with unvoiced/voiced consonants^9^, which are characterized by distinctive spectral profiles. Unvoiced consonants (/k/, /t/) are characterized by stronger energy in higher frequencies and show broader, abrupt and noisier distribution of power whereas voiced consonants (/b/, /d/) elicit energy in lower frequencies arising from the regular pulses of voicing during phonation. These spectral characteristics are represented in the spectro-temporal distribution (FFT) and the overall distribution of spectral power (spectral tilt). Vocal roughness has been shown to exaggerate the apparent body size of the vocalizer^67^. Since the proportion of unvoiced segments is indicative of vocal roughness, the fraction of unvoiced frames is an important parameter. Previously, Knoeferle et al.^23^ showed that the duration of vowels in pseudowords influences sound-size judgements, however, with the current set of pseudowords, we found that bigness, but not smallness, ratings were associated with the back rounded vowels^9^. Thus, pseudoword duration, which is partly contingent on the component vowels, is relevant for size ratings. F0 and parameters representing frequency variability, e.g. jitter and F0_SD_; amplitude variability (shimmer); periodicity, e.g. HNR, pulse number and mean autocorrelation; did not feature in the combination.

### 3. Brightness

a. *RSA for spectro-temporal parameters*

RSA revealed weak but statistically significant correlations between the ratings RDM and all three spectro-temporal parameter RDMs (**Figure 5B-D**). The strongest among them was for spectral tilt (r = .14, p < .0001), followed by weaker correlations for the FFT (r = .08, p < .0001) and the speech envelope (r = .04, p < .0001). This pattern suggests that the distribution of spectral energy is more important than temporal modulation to distinguish bright- and dark-rated pseudowords.

**Figure 5:**
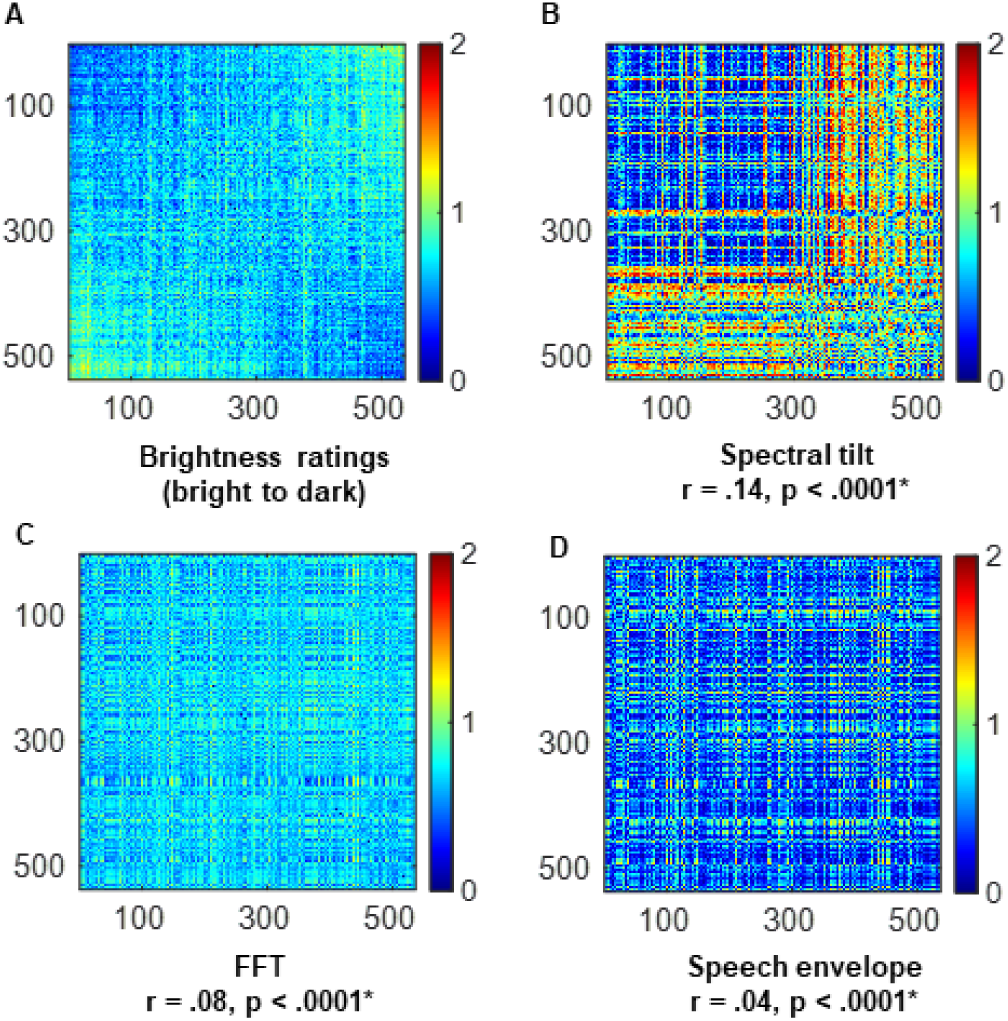
Representational similarity analyses and conventional correlations relating auditory brightness ratings and acoustic parameters. **A**: Ratings RDM. **B–D**: RDMs for the three spectro-temporal parameters. Pseudowords are ordered, left to right, from brightest to darkest. All other details as for Fig. 3.

Spectral tilt was flatter for brighter-rated pseudowords, reflecting more high-frequency energy, whereas for darker-rated pseudowords, power was concentrated in the lower frequency bands (**Supplementary** Figure 13a). These differences are consistent with the phonetic composition of the stimuli, where pseudowords rated as bright more often comprise unvoiced consonants and front unrounded vowels, while pseudowords rated as dark tend to feature voiced consonants and back rounded vowels. The spectrogram showed greater power in the higher frequencies for pseudowords rated as bright, compared to those rated as dark (**Supplementary** Figure 13b). The speech envelope was discontinuous for both bright- and dark-rated pseudowords but amplitude was perhaps greater for the latter (**Supplementary** Figure 13c), reflecting that bright/dark-rated pseudowords tend to involve unvoiced/voiced consonants, respectively.

*Conventional correlations for voice parameters*

We observed that the fraction of unvoiced frames decreased significantly as ratings increased towards the dark end of the scale (r = -.20, p < .001), while mean HNR (r = .20, p < .001) and pulse number (r = .13, p = .004) were significantly positively correlated with ratings such that voice quality became more periodic and smoother for pseudowords that were rated as darker, indicating that darker-rated pseudowords are produced with greater periodicity and a higher proportion of voiced segments^9^. F0 decreased significantly as ratings shifted towards the dark end of the scale (r = -.17, p < .001). This aligns with prior findings linking high pitch to brightness and low pitch to darkness, as brighter-rated pseudowords tended to exhibit higher F0, while darker-rated pseudowords were characterized by lower F0 and stronger low-frequency energy^7,9,68^. In line with Tzeng et al.^69^, there was a small effect of duration (r = .1, p = 02) in which longer pseudowords were rated as darker, but this did not survive correction. There were also weak relationships for shimmer (r = -.11, p = .01) and MAC (r = .1, p = .02) that did not survive correction, and non-significant relationships were found for F0_SD_ (r = -.05, p = .2) and jitter (r = -.04, p = .4) (**Table 1** and **Supplementary** Figure 14a-i). Together, these results suggest that brightness and darkness judgments are primarily guided by spectral distribution and voicing-related periodicity rather than by fine-grained variability in amplitude or frequency.

*Optimal combination*

A combination of ten parameters including all three spectro-temporal parameters: FFT, spectral tilt, and speech envelope; and seven voice parameters: fraction of unvoiced frames, pulse number, jitter, MAC, F0, F0_SD_ and duration, constituted the optimal model that showed the best correlation (ρ = 0.52) between perceptual brightness ratings (N = 50) and those generated by the KNN algorithm (**Table 2**). Lacey et al.^9^ found that, similar to the size domain, consonant voicing and place of articulation were key determinants of the brightness ratings of pseudowords. These characteristics change the frequency profiles of the speech signal by shifting energy toward lower or higher bands and are therefore captured by the FFT, explaining its presence in the combination. Additionally, consonant voicing and vowel category affect spectral tilt, e.g., voiced consonants and back rounded vowels have more energy in the lower frequencies, while unvoiced consonants and front unrounded vowels have more energy in the higher frequencies,^20^ explaining the importance of spectral tilt in the combination. The temporal evolution of pseudowords also reliably captures information related to vowel rounding, which influences brightness ratings^9^. This is represented by the speech envelope, which explains its presence in the model. Furthermore, information related to voicing, represented by the fraction of unvoiced frames, is crucial because the key difference between pseudowords rated as bright vs. dark is the presence of unvoiced vs. voiced consonants^9^. The contribution of F0 and F0_SD_ is consistent with the finding that participants match high-pitched sounds to lighter, and low-pitched sounds to darker, stimuli^70^. As for the shape domain, pseudowords rated as bright/dark were characterized by the presence of unrounded/rounded vowels, which influence the duration of pseudowords, thus explaining the relevance of duration in brightness judgements. The inclusion of MAC, pulse number and jitter reflect that measures of periodicity and stability of vocal fold vibration mediate sound-brightness judgements. Pulse number is influenced by phonetic properties such as consonant voicing and vowel category: voiced consonants and vowels involve regular vocal fold vibration, resulting in more glottal pulses, whereas voiceless consonants introduce aperiodic segments with fewer pulses. Jitter and mean autocorrelation capture regularity in the waveform; greater regularity (high mean autocorrelation and low jitter) is associated with voiced and sonorant segments, while irregular or absent periodicity reflects unvoiced segments. Measures like jitter and MAC robustly quantify voicing stability^66,67^, and are sensitive to the presence/absence of vocal fold vibration across segments. Thus, their inclusion provides cues that distinguish voiced vs. unvoiced contrasts that appear to be central to brightness judgments. The parameter related to amplitude variability (shimmer) did not feature in the combination for brightness; neither did the HNR, although other measures of periodicity (see above) did.

Again, there were some discrepancies between the parameters found to be significant on the analyses of individual parameter relationships to ratings, and those featuring in the optimal model. Thus, the fraction of unvoiced frames and the HNR were significantly correlated with ratings individually, but were not part of the optimal combination in the KNN approach. Conversely, the KNN model included parameters (MAC, jitter, F0, F0_SD_ and duration) that were not deemed significant on the individual analyses.

### 4. Hardness

a. *RSA for spectro-temporal parameters*

The RDMs for all three spectro-temporal parameters were significantly positively correlated with the RDM for ratings of the pseudowords from hard to soft (**Figure 6B-D**). The FFT RDM showed the strongest correlation (r = .36, p < .0001). The RDM of the speech envelope was similarly strongly correlated (r = .33, p < .0001) while the relationship between ratings and spectral tilt was weaker (r = .11, p < .0001). Hard-rated pseudowords comprised obstruents produced at alveolar and post-alveolar or velar places of articulation, which generate bursts of high-frequency energy and sharp changes in amplitude. In contrast, soft pseudowords were more often composed of sonorants and bilabial or labiodental articulations, as well as back rounded vowels, producing smoother and more continuous acoustic patterns^9^. These differences are consistent with the presence of greater high-frequency energy and more abrupt spectro-temporal transitions in pseudowords judged as hard, compared to the more gradual and lower-frequency variations found in pseudowords considered soft (**Supplementary** Figure 15a). The spectral tilt results align with this interpretation, as hard-rated pseudowords exhibited a flatter tilt reflecting greater energy at higher frequencies, whereas soft-rated pseudowords showed a steeper tilt consistent with reduced high-frequency energy (**Supplementary** Figure 15b). The speech envelope further reflects these phonetic contrasts, i.e., pseudowords rated as hard displayed more pronounced discontinuities and irregular amplitude fluctuations, while pseudowords rated as soft showed smoother and more continuous envelopes (**Supplementary** Figure 15c). These contrasts underlie the observed correlation between the speech envelope and perceptual ratings, suggesting that the temporal dynamics of the acoustic signal contribute to sound-hardness associations.

*Conventional correlations for voice parameters*

**Figure 6.**
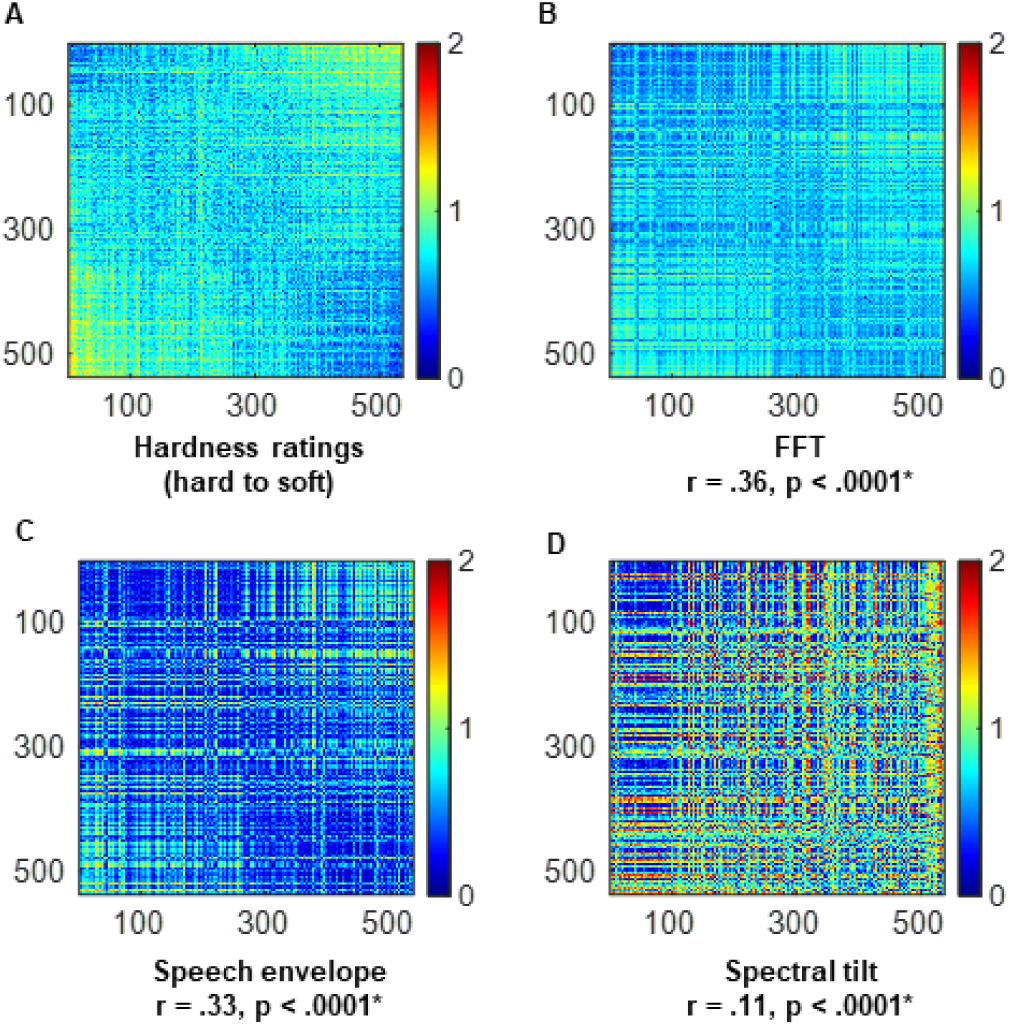
Representational similarity analyses and conventional correlations relating auditory hardness ratings and acoustic parameters. **A**: Ratings RDM. **B–D**: RDMs for the three spectro-temporal parameters. Pseudowords are ordered, left to right, from hardest to softest. All other details as for Fig. 3

For the voice parameters, soft-rated pseudowords were associated with greater periodicity and less vocal variability, while the reverse was true for the hard-rated pseudowords (**Table 1** and **Supplementary** Figure 16a-i). Thus, the mean HNR (r = .42, p < .001), pulse number (r = .38, p < .001), and MAC (r = .30, p < .001) were significantly positively correlated with ratings, while the fraction of unvoiced frames (r = -.28, p < .001) was significantly negatively correlated. These patterns indicate that higher perceived softness is characterized by more periodic and acoustically regular vocal signals, consistent with greater smoothness and continuity in phonation. Shimmer (r = -.26, p < .001) and jitter (r = -.23, p < .001) were significantly negatively correlated with hardness ratings, meaning that as ratings moved from hard to soft, there was decreasing variability in amplitude and frequency, respectively. This reduction in vocal instability suggests that softness is conveyed by acoustic regularity and low perturbation, whereas hardness is cued by aperiodicity and roughness in the vocal signal. The sharper spectral transitions and high-frequency emphasis of hard-rated pseudowords are consistent with the presence of obstruents, particularly fricatives, which are noisy and aperiodic, whereas the smoother, more periodic profiles of soft-rated pseudowords align with sonorant-like articulation. Duration (r = .30, p < .001) was positively related to softness, aligning with underlying phonetic contrasts in obstruent versus sonorant content for hard- and soft-rated pseudowords. Hardness ratings were uncorrelated with F0 (r = - 0.01, p = 0.7); there was a weak effect in which F0_SD_ (r = -0.12, p = 0.006) decreased for softer pseudowords but this did not survive correction. Thus, the fundamental frequency and its standard deviation do not appear to provide consistent cues mediating iconicity judgments of hardness.

*Optimal combination*

The optimal combination that yielded the highest correlation between the KNN-generated ratings and participants’ perceptual ratings (N = 54) of hardness comprised five acoustic parameters (ρ = 0.54). This set included two spectro-temporal parameters: FFT and spectral tilt, and three voice parameters: fraction of unvoiced frames, shimmer, and duration (**Table 2**). The place of articulation of sonorants significantly predicts softness ratings, while voicing and manner and place of articulation are key predictors of hardness ratings^9,71,72^. These features have very distinct spectro-temporal profiles, effectively captured by the FFT, explaining the importance of this parameter in the combination. While these finer details are crucial for sound-hardness associations, hard- and soft-rated pseudowords can also be distinguished by the broad phonetic categories of obstruents and sonorants^9^, respectively, which are reflected in spectral tilt. Consequently, spectral tilt is also influential. Furthermore, the utterance of an obstruent is characterized by discontinuous and turbulent airflow with major constrictions to the vocal tract, and the fraction of unvoiced frames reflects such changes. The contributions of shimmer and duration are consistent with prior studies showing that noisy and punctuated sounds with abrupt and highly variable onsets, involving voiceless plosives (e.g. /t/ and /k/) and bilabial plosives (e.g. /p/ and /b/) are associated with the tactile perception of softness^72^. Such phonemes are characterized by distinctive temporal variations in amplitude, measured by shimmer, and by durational cues. As for other domains, parameters that indicate frequency variability, e.g. jitter and F0_SD_; the amplitude profile, i.e. the speech envelope; and periodicity, e.g. HNR, pulse number and MAC; and F0 did not contribute to predicting hardness ratings.

As for the other domains above, the different analytical approaches produced some discrepancies for the hardness domain: the speech envelope, periodicity measures (HNR, pulse number, MAC), F0, and F0_SD_ were not included in the optimal KNN model despite being individually significant; however, the optimal combination only comprised factors that were found to be significant on the individual analyses.

### 5. Roughness

a. *RSA for spectro-temporal parameters*

RSA showed that the RDMs for all three spectro-temporal parameters were significantly positively correlated with the RDM for ratings of the pseudowords from smooth to rough (**Figure 7B–D**). The strongest correlation was for the FFT RDM (r = .32, p < .0001), followed by the speech envelope RDM (r = .28, p < .0001). Spectral tilt had the weakest relationship to the ratings in terms of between-RDM correlations (r = .10, p < .0001). This pattern suggests that both spectral composition and amplitude modulation provide reliable cues for perceived roughness, with FFT and the speech envelope most closely reflecting how listeners map sound to smoothness or roughness. Pseudowords rated as smooth were characterized by more gradual transitions in energy across frequencies and time (**Supplementary** Figure 17a) compared to those rated rough, which had more abrupt transitions. These smoother patterns likely arise from sonorant-dominated segments and voiced consonants that involve open and stable vocal tract configurations, producing harmonic-rich and continuous energy distributions across time and frequency^71^. In contrast, rough- rated pseudowords contain more voiceless obstruents such as stops and fricatives^73,74^, which involve turbulent airflow and abrupt closures that yield aperiodic and discontinuous energy patterns. The speech envelope was more even and continuous for smooth-rated pseudowords but uneven and discontinuous for rough-rated pseudowords (**Supplementary** Figure 17b). This distinction mirrors articulatory differences between sustained, resonant voicing and transient, noise-like gestures. More power was concentrated in higher frequencies for smooth-rated compared to rough-rated pseudowords (**Supplementary** Figure 17c). This higher-frequency emphasis in smooth items is consistent with the formant structure of sonorants and voiced segments, whereas the broadband, noisier spectra of rough items reflect the turbulent properties of voiceless obstruents^75^.

*Conventional correlations for voice parameters*

**Figure 7:**
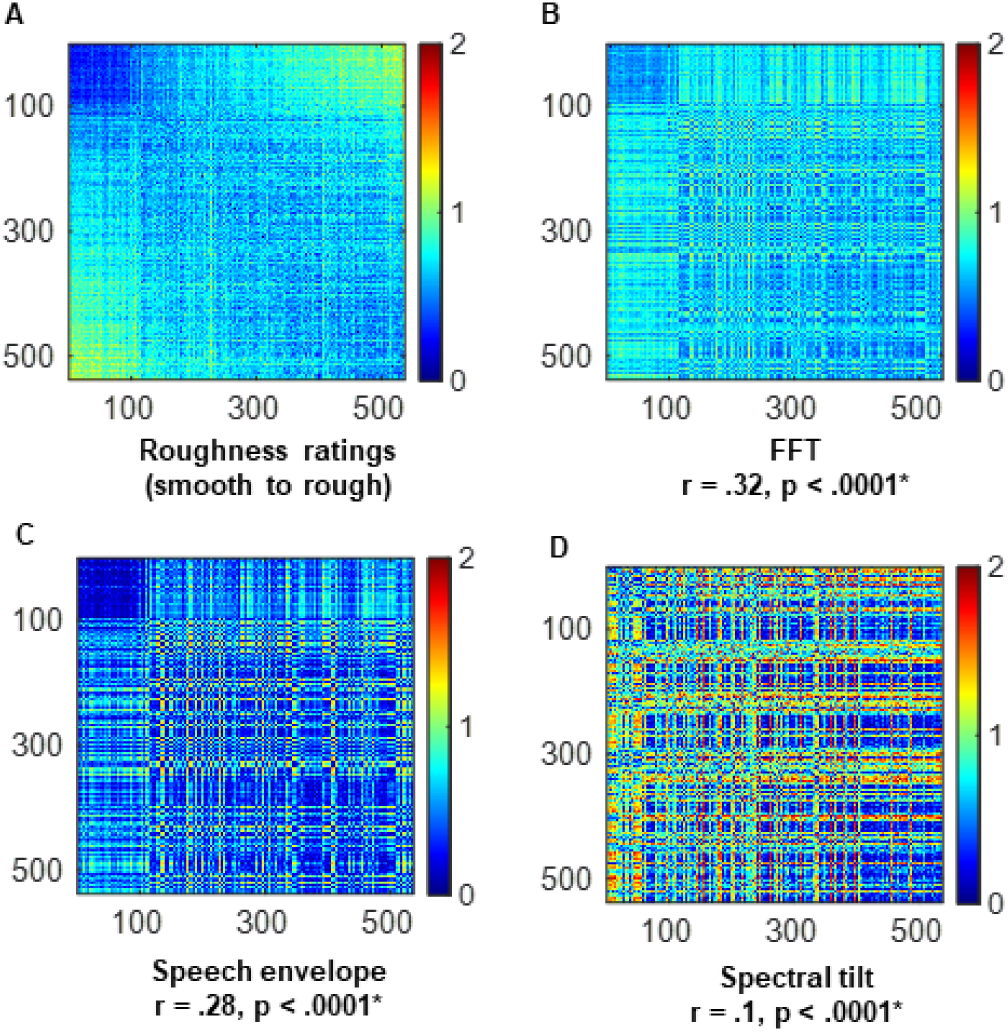
Representational similarity analyses and conventional correlations relating auditory roughness ratings and acoustic parameters. **A**: Ratings RDM. **B–D**: RDMs for the three spectro-temporal parameters. Pseudowords are ordered, left to right, from smoothest to roughest. All other details as for Fig. 3

For the voice parameters (**Table 1** **and Supplementary** Figure 18a–i), the HNR (r = –.51, p < .001), pulse number (r = –.40, p < .001), MAC (r = –.36, p < .001), and duration (r = –.20, p < .001) were significantly negatively correlated with ratings of the pseudowords, indicating that periodicity and duration decreased as ratings transitioned from smooth to rough. This suggests that smoother-rated pseudowords are produced with more periodic and stable voicing, whereas rougher-rated ones involve a higher proportion of unvoiced or irregular segments that disrupt vocal periodicity. The decline of pseudoword duration towards the rough end suggests that shorter and more temporally compact pseudowords are perceived as rougher, consistent with previous findings that rapid, abrupt gestures cue tactile-like roughness^76^. The fraction of unvoiced frames (FUF; r = .37, p < .001) increased from smooth to rough, reflecting a rise in aperiodic and turbulent segments. Similarly, shimmer (r = .33, p < .001) and jitter (r = .23, p < .001) were significantly positively correlated with roughness ratings, indicating greater variability in amplitude and frequency as pseudowords were perceived as rougher. This variability likely reflects the articulatory turbulence and unstable vocal fold vibration typical of voiceless consonants, whereas smoother items are characterized by more regular and consistent phonation. The mean F0 (r = .02, p = .51) and its variability (F0_SD_; r = .10, p = .01) were not reliably correlated with ratings, indicating that pitch and its fluctuations contribute little to perceived smoothness or roughness.

*Optimal combination*

The KNN algorithm identified an optimal combination comprising the two spectro-temporal parameters: FFT and speech envelope, that produced ratings for the 537 pseudowords which correlated (ρ =.56) best with the perceptual ratings (N = 58) of pseudowords in the roughness domain (**Table 2**). Prior research indicates that, as noted above, items rated as smooth are dominated by sonorants and voiced segments, comprising gradual formant transitions and sustained amplitude, the features well captured by FFT and speech envelope. In contrast, items rated as rough are characterized by voiceless obstruents that show broadband noise bursts and abrupt energy changes, resulting in jagged, discontinuous envelopes. Because roughness perception depends on the balance between gradual/abrupt spectral energy and temporal continuity, the combination of information represented in the FFT and speech-envelope carries cues that mediate iconicity judgements of roughness.

The optimal KNN model comprised only 2 of 10 parameters that were individually found to be significant: spectral tilt, measures of periodicity (HNR, pulse number, MAC), voicing and its variability (fraction of unvoiced frames, shimmer, jitter, F0_SD_), F0, and duration were all significant on individual analyses but did not feature in the optimal model.

### 6. Weight

a. *RSA for spectro-temporal parameters*

RSA showed that the RDMs for all three spectro-temporal parameters were significantly positively correlated with the RDM for ratings of the pseudowords from light to heavy (**Figure 8B–D**). The strongest correlation was for the FFT RDM (r = .38, p < .0001), followed by the speech envelope RDM (r = .28, p < .0001), while spectral tilt showed the weakest relationship (r = .19, p < .0001). This indicates that both the spectral structure and temporal amplitude profile of pseudowords contribute to perceived weight, with FFT capturing the most informative cues for distinguishing lighter from heavier sounds. Pseudowords rated as light exhibited smoother, continuous energy transitions across frequency and time, whereas those rated as heavy showed sharper and more discontinuous transitions (**Supplementary** Figure 19a**, b**). These acoustic patterns likely reflect underlying phonetic distinctions, where lighter-rated pseudowords tend to include sonorants and labial or labiodental articulations with unrounded vowels, while those rated as heavier contain voiced obstruents produced at post-alveolar or velar places of articulation, and rounded vowels^9^. The distribution of these articulatory features accounts for the spectro-temporal profiles: sonorants produce steady, harmonic-rich energy, whereas obstruents create irregular and transient bursts. The relatively weak association of spectral tilt with perceptual ratings may result from the mixed presence of low- and high-frequency energy in both light- and heavy-rated pseudowords (**Supplementary** Figure 19c).

*Conventional correlations for voice parameters*

**Figure 8:**
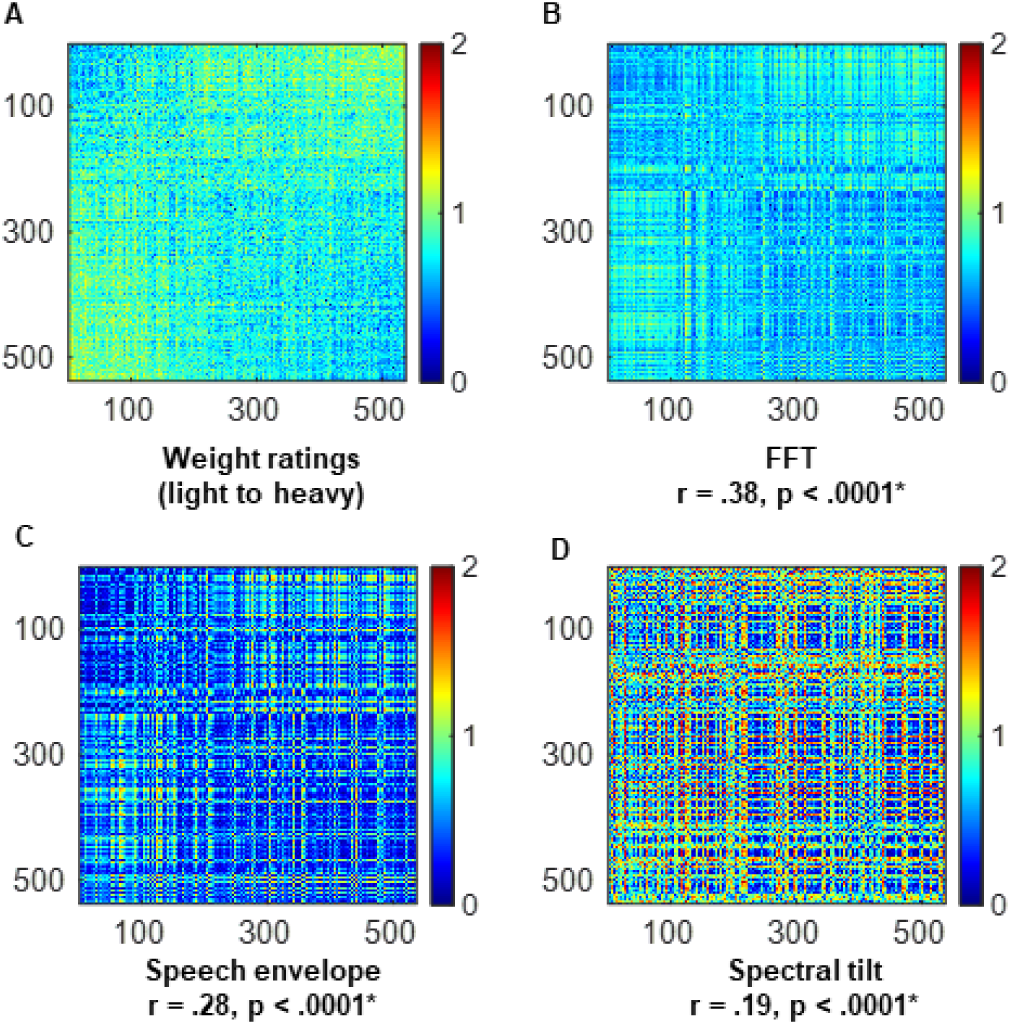
Representational similarity analyses and conventional correlations relating auditory ratings of weight and acoustic parameters. **A**: Ratings RDM. **B–D**: RDMs for the three spectro-temporal parameters. Pseudowords are ordered, left to right, from lightest to heaviest. All other details as for Fig. 3

Among the voice parameters (**Table 1** **and Supplementary** Figure 20a–i), the HNR (r = –.34, p < .001), pulse number (r = –.29, p < .001), and MAC (r = –.28, p < .001) were significantly negatively correlated with ratings, indicating that periodicity decreased as ratings transitioned from light to heavy. This reduction in periodicity suggests a shift from voiced to unvoiced segments, consistent with a broader transition from sonorants to obstruents as perceived weight increases^9^. The fraction of unvoiced frames (r = .16, p < .001) increased as ratings transitioned from light to heavy, fitting with greater aperiodicity in heavy-rated pseudowords. Similarly, shimmer (r = .21, p < .001), jitter (r = .20, p < .001) and F0_SD_ (r = .15, p < .001) were positively correlated with heaviness, indicating that more variable voicing contributes to the perception of heaviness, possibly due to increased articulatory effort or instability in vocal fold vibration. F0 (r = –.14, p < .001) was lower for heavy-rated pseudowords than for light-rated ones, mirroring natural size– weight correspondences where higher pitch is associated with smaller and lighter objects and lower pitch with larger and heavier ones. Duration (r = –.20, p < .001) was also negatively correlated with ratings, suggesting that shorter, more temporally compact pseudowords are perceived as denser and therefore heavier, a relationship that warrants further investigation in direct tests of perceived density.

*Optimal combination*

The KNN algorithm identified a combination of six parameters including three spectro-temporal parameters: FFT, spectral tilt, and speech envelope, and three voice-related parameters: fraction of unvoiced frames, HNR, and duration, that generated predicted ratings for the 537 pseudowords with the strongest correlation (ρ = 0.45) to the perceptual ratings provided by participants (N = 55) in the weight domain (**Table 2**). Lacey et al.^9^ showed that the combinations of vowel rounding and the consonant features that influence light and heavy ratings are unique. For example, light ratings are influenced by the presence of unrounded vowels along with unvoiced bilabial and labiodental stops, while heavy ratings were reliant on back rounded vowels along with voiced post alveolar/velar stops and a/fricatives. Fricative sounds (e.g., /f/) involve turbulent airflow, resulting in lower HNR. In contrast, voiced consonant sounds (e.g., alveolar: /d/ and velar /g/) have a more periodic nature and less turbulent airflow. Since the HNR indicates the proportion of periodic and aperiodic components in speech, it significantly influences weight ratings of pseudowords. Sonorants and obstruents that determine the lightness and heaviness of the pseudowords, respectively, show distinct spectral tilts, accounting for the influence of spectral tilt. A characteristic distinction between these two classes of phonemes (i.e., sonorants and obstruents) is also their voicing. The FUF captures sensitivity to voicing, while the speech envelope reflects differences between obstruents and sonorants through amplitude. Overall, voiced segment proportion and amplitude strongly influenced weight judgments, while duration likely reflected vowel effects. Other parameters including frequency and amplitude variability, periodicity measures, and F0 did not predict weight ratings.

While all 12 parameters were found to be significant individually, the optimal model consisted of only 6 of them: the 3 spectro-temporal ones, the HNR, fraction of unvoiced frames, and duration. Parameters denoting variability of voicing: shimmer, jitter, and F0_SD_; F0, and some parameters of periodicity: pulse number and MAC; were omitted from the model.

### 7. Arousal

a. *RSA for spectro-temporal parameters*

Significant positive correlations with the ratings RDM were found for all three spectro-temporal parameter RDMs (**Figure 9B–D**), although they were weaker than those observed for the shape,

*Conventional correlations for voice parameters*

**Figure 9:**
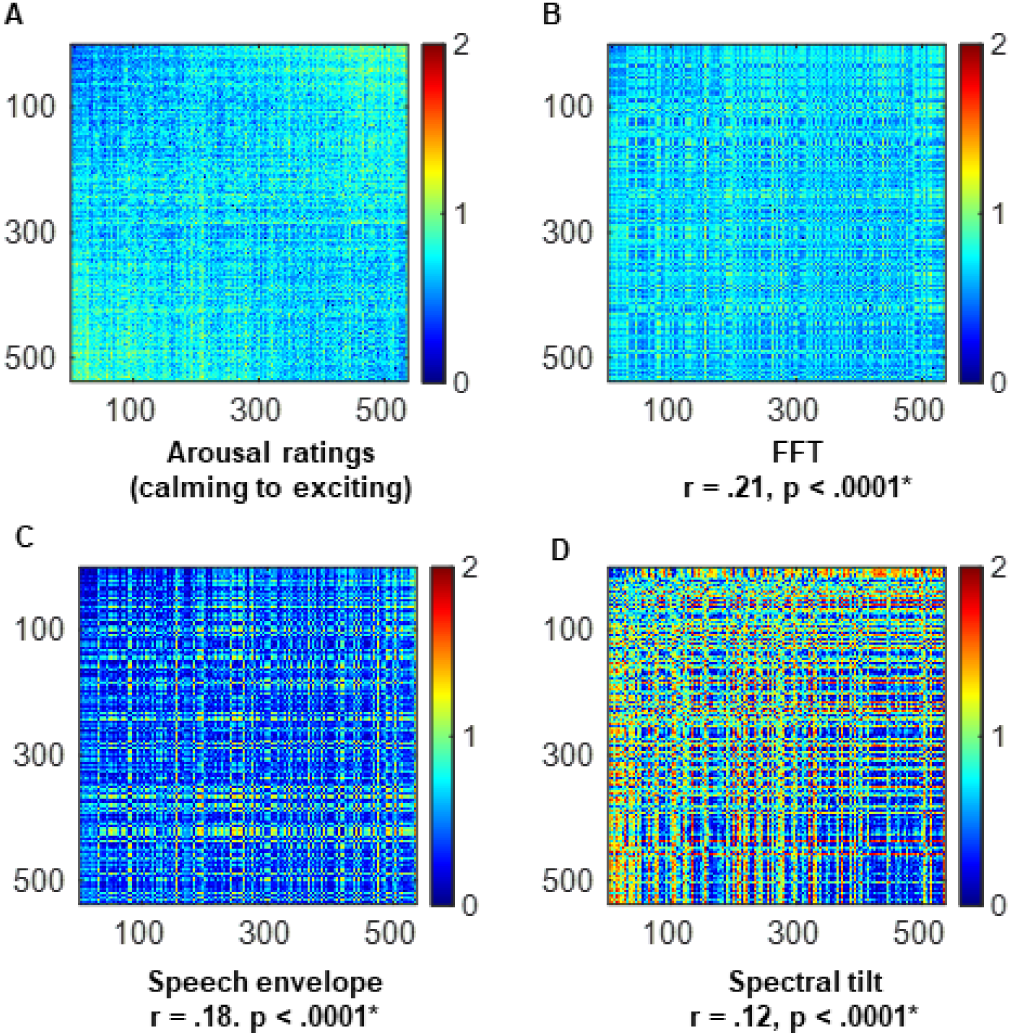
Representational similarity analyses and conventional correlations relating auditory ratings of arousal and acoustic parameters. **A**: Ratings RDM. **B–D**: RDMs for the three spectro-temporal parameters. Pseudowords are ordered, left to right, from calming to exciting. All other details as for Fig. 3 weight, and hardness domains. The highest correlation with the arousal ratings RDM was for the FFT RDM (r = .21, p < .0001), followed by the speech envelope RDM (r = .18, p < .0001), and then the spectral tilt RDM (r = .12, p < .0001). This pattern indicates that both spectral and temporal dynamics contribute to perceived emotional arousal, with FFT capturing the most salient cues distinguishing calming- from exciting-rated pseudowords. Changes in the distribution of power across frequencies over time were more gradual for calming-rated pseudowords and more abrupt for those rated as exciting (**Supplementary** Figure 21a). These acoustic differences likely arise from phonetic contrasts, where pseudowords perceived as calming tend to contain sonorants and back rounded vowels, while those perceived as exciting more often include unvoiced obstruents and front unrounded vowels^9,74^. For calming-rated items, the speech envelope was relatively continuous and the spectral tilt was steep, reflecting smoother transitions and stronger energy in lower frequencies associated with bilabial or labiodental articulation. This configuration aligns with the sustained amplitude and low-frequency emphasis characteristic of sonorants and bilabials. In contrast, pseudowords considered as exciting displayed more discontinuous envelopes and flatter spectral tilt, with energy distributed across frequencies (**Supplementary** Figure 21b–c). These properties are consistent with rapid articulatory gestures and turbulent airflow in stops and fricatives, which produce irregular temporal patterns and high-frequency energy.

The voice parameters HNR (r = -.42, p < .001), pulse number (r = -.30, p < .001), and MAC (r = - .26, p < .001) were negatively correlated with ratings, indicating that periodicity decreased as ratings transitioned from calming to exciting (**Table 1** and **Supplementary** Figure 22a-i). *This reduction in periodicity suggests that exciting-rated pseudowords are produced with less stable voicing and greater vocal effort, consistent with increased physiological arousal*^9,77^. The fraction of unvoiced frames was positively correlated (r = .40, p < .001) with ratings, meaning that the number of unvoiced segments increased from calming to exciting, reinforcing the pattern of greater aperiodicity and turbulence. Jitter (r = .14, p < .001) and shimmer (r = .29, p < .001) also rose with arousal, reflecting heightened variability in amplitude and frequency linked to expressive or tense phonation. In contrast, F0 (r = .07, p = .10), F0_SD_ (r = .07, p = .08), and duration (r = .02, p = .50) showed no significant relationships with ratings, suggesting that pitch and timing contribute less to the perception of emotional arousal than do spectral and voicing cues.

*Optimal combination*

The optimal combination that produced ratings correlating best (ρ = 0.51, N = 58) with participants’ perceptual ratings of arousal comprised five acoustic parameters, including the three spectro-temporal parameters: FFT, spectral tilt and speech envelope; and two voice parameters: the fraction of unvoiced frames, and HNR (**Table 2**). Pseudowords rated as calming comprise sonorants and back rounded vowels while those rated as exciting primarily comprise unvoiced obstruents and front unrounded vowels^9,71^. Spectro-temporal features that differentiate sonorants from obstruents, as well as those related to voicing, are effectively captured by the FFT as well as the speech envelope. Additionally, spectral tilt captures the broad distinction in the spectral profile of sonorants and obstruents. This explains the presence of the spectro-temporal parameters in the optimal combination. Affective states induced by stress have been shown to influence the HNR in speech produced during these states^78^. Additionally, the presence of obstruents increases turbulent airflow at the glottis during phonation, leading to aperiodic vocal fold vibration, which impacts the HNR. Also, voicing of obstruents underlie the tendency for pseudowords to be rated as exciting, explaining the significance of the FUF in the model.

The optimal combination according to the KNN approach included 5 of the 9 parameters found to be significant on individual analyses: the 3 spectro-temporal ones, the HNR, and the fraction of unvoiced frames; whereas the other individually significant parameters (pulse number, MAC, shimmer, jitter) were absent from the optimal model.

### 8. Valence

a. *RSA for spectro-temporal parameters*

Weak, but significant, positive correlations were found for all three spectro-temporal parameter RDMs with the ratings RDM (**Figure 10B-D**). The FFT RDM had the highest correlation of the three (r = .17, p < .0001), followed by the speech envelope RDM (r = .16, p < .0001) and then the spectral tilt RDM (r = .07, p < .0001). This pattern indicates that both spectral and temporal information contribute modestly to valence perception, with FFT carrying more informative cues for distinguishing good-from bad-rated pseudowords. The spectrograms showed changes in the distribution of power across frequency over time that were more diffuse for pseudowords rated as good, compared to those rated as bad (**Supplementary** Figure 23a). These acoustic trends likely arise from phonetic differences: good-rated pseudowords, containing a higher proportion of sonorants and unrounded front vowels, produced smoother and more continuous spectral trajectories, while bad-rated pseudowords, which contained more obstruents and rounded back vowels, produced sharper and more discontinuous changes in energy distribution^9^. The speech envelope was smoother for good-rated pseudowords and more discontinuous for bad-rated pseudowords, consistent with differences in phonemic amplitude and timing that accompany the contrast between sonorants and obstruents. (**Supplementary** Figure 23b). The minimal effect of spectral tilt suggests that both categories shared comparable distributions of high- and low-frequency energy, leading to little variation in slope across the valence continuum. (**Supplementary** Figure 23c).

*Conventional correlations for voice parameters*

**Figure 10:**
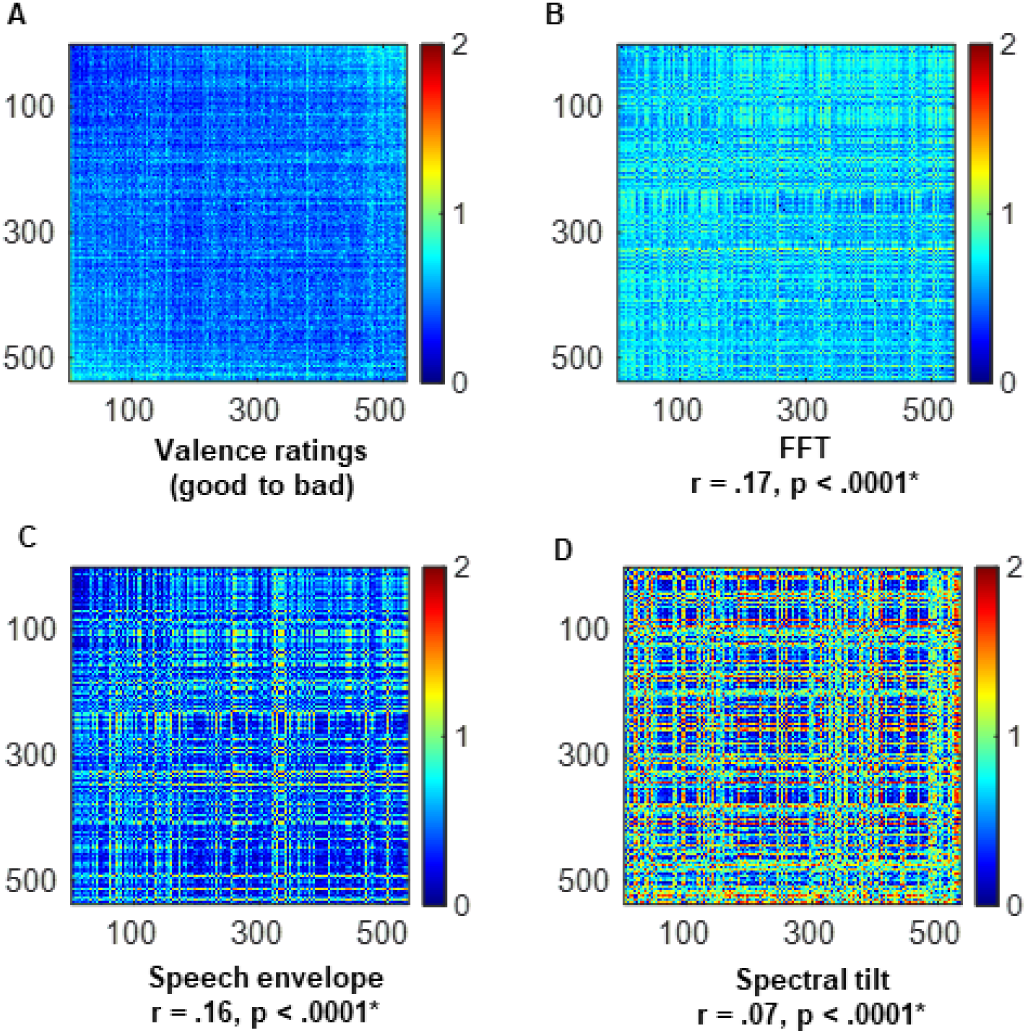
Representational similarity analyses and conventional correlations relating auditory ratings of valence and acoustic parameters. **A**: Ratings RDM. **B–D**: RDMs for the three spectro-temporal parameters. Pseudowords are ordered, left to right, from good to bad. All other details as for Fig. 3

Among the voice parameters, the mean HNR (r = -.40, p < .001), pulse number (r = -.37, p < .001), and MAC (r = -.35, p < .001) were significantly negatively correlated with ratings, and the fraction of unvoiced frames (r = .36, p < .001) was significantly positively correlated with ratings. This pattern reflects decreasing periodicity and increasing aperiodicity as pseudowords shift from good to bad, suggesting greater irregularity and instability in voicing for items perceived to be bad. Shimmer (r = .36, p < .001), jitter (r = .26, p < .001), and F0_SD_ (r = .23, p < .001) were significantly positively correlated with ratings, reflecting increasing variability in voice quality for pseudowords perceived as bad, compared to those perceived as good. A significant negative correlation with ratings indicated that F0 (r = -.20, p < .001) was lower for bad-rated, compared to good-rated, pseudowords. Duration (r = .16, p < .001) was not significantly correlated with the valence ratings (**Table 1** and **Supplementary** Figure 24a-i). Together, these results suggest that listeners’ judgments of pseudoword valence are shaped by consistent acoustic and phonetic cues, where smoother, more periodic, and harmonically rich vocal patterns correspond to positive impressions, and rougher, aperiodic, and variable patterns correspond to negative ones.

*Optimal combination*

The KNN algorithm identified an optimal combination comprising the two spectro-temporal parameters: FFT and speech envelope, that produced ratings for the 537 pseudowords which correlated (ρ =.66) best with the perceptual ratings (N = 49) of pseudowords in the valence domain (**Table 2).** The primary characteristic distinguishing the pseudowords representing the categorical opposites in the valence domain is the incidence of sonorants/obstruents^9^. Spectro-temporal features that differentiate sonorants from obstruents, as well as those related to voicing, are effectively captured by the FFT as well as the speech envelope. Parameters that feature frequency variability (spectral tilt, jitter, F0_SD_) amplitude variability (shimmer); periodicity (the fraction of unvoiced frames, pulse number and mean autocorrelation) and F0, did not contribute to predicting the arousal ratings in the optimal KNN model, despite being individually significant. Duration was neither part of the optimal model nor was it individually significant.

### **B.** Predicting iconicity shape ratings of real words

We used the multidimensional space defined by the five acoustic parameters that best predicted the shape ratings of the pseudowords (see section III.A.1) to predict the shape ratings of real words. The predicted rating of each real word was based on the modal perceptual rating of the 23 pseudowords in its acoustic neighborhood. Since the pseudowords were rated by 60 participants, the mean across-participant pseudoword ratings were used to predict ratings of the real words. The predicted ratings were strongly correlated (*r* = 0.52, 𝑝 < 0.001) with mean perceptual ratings of the real words.

## **IV.** DISCUSSION

Behavioral^3–5^, developmental^11,79^, and neuroimaging^80,81^ studies have widely employed pseudowords to demonstrate the phenomenon of sound iconicity and to study the cognitive processes that underlie iconic associations of sounds. The acoustic parameters that underpin sound iconicity have been less well studied. Prior studies investigating acoustic properties that mediate iconicity suggest that the influence of these properties varies by semantic domain^23,29^. However, the associations of acoustic properties with iconic mappings have not previously been investigated systematically across a range of semantic domains. Here, building on a prior study from our group^29^, we assessed the relationship between each of twelve acoustic parameters and sound iconicity ratings in eight different meaning domains, using RSA for the three spectro-temporal parameters studied and conventional correlations for nine parameters associated with voice quality.

Additionally, we identified the combinations of acoustic parameters that correlate best with iconic associations in each domain. We reasoned that if a set of iconic pseudowords were projected onto a multidimensional acoustic space, defined by the optimal combination of acoustic parameters for a particular domain of meaning, then pseudowords in that space would tend to form coherent clusters, i.e., pseudowords with similar degrees of sound-iconic associations would be close to each other. Based on this premise, we employed a supervised KNN machine-learning approach and found that domain-specific combinations of acoustic parameters yielded predicted ratings that correlated with the perceptual ratings of pseudowords across the eight different meaning domains. Finally, as a proof of concept, we used the KNN approach to generate iconicity shape ratings of real words based entirely on their acoustic properties, minimizing semantic bias.

We chose the KNN algorithm because it is well suited to our idea that pseudowords with similar sound-iconic ratings cluster together in a multidimensional acoustic parameter space. Unlike approaches relying on methods such as support vector machines (SVM), regression, or deep learning, which require fitting a global decision boundary, a parametric mapping, or a complex transformation of the input features, KNN operates directly on the local geometry of the parameter space. This makes it easily interpretable in the context of our goal: if two pseudowords are acoustically similar, they should be close in the parameter space and therefore have similar ratings, and our KNN based algorithm captures this relationship. Moreover, KNN’s distance-based metric allows us to quantify acoustic proximity, which in turn reflects perceptual similarity. While SVM and regression models can perform well for prediction, they both tend to be more or less opaque regarding their underlying neighborhood structure^82^; and deep learning methods, though powerful, would require substantially larger datasets, risk overfitting, and make interpretability of specific parameter contributions more challenging. In contrast, KNN allows us to interpret the multidimensional clusters of pseudowords, making it a theoretically and methodologically appropriate choice for our study.

### Complementary insights from multiple analyses

As noted above, we observed discrepancies between our analyses of the relationships between individual parameters and ratings (using RSA and conventional correlations), and the optimal combinations of features that emerged in our KNN models. These mismatches highlight the varying sensitivities of the three analytical frameworks. Conventional correlations capture overall, linear correspondences between individual parameters and perceptual ratings, while RSA can reveal more granular relationships over a time and/or frequency axis. These analyses inform how individual parameters vary systematically with iconicity ratings. However, the KNN models are based on the local geometry within a multidimensional parameter space. Parameters that show only small relationships on an individual basis can nonetheless reshape the local topology of this space. On the other hand, features with strong individual correlations may contribute redundant variance, offering limited new information once other correlated or complementary cues are included. Overall, our findings reveal that iconicity associations emerge not from isolated acoustic cues but from a combination of interacting parameters.

### Common and differing acoustic cues across meaning domains

We found that the best-performing combinations of acoustic parameters for different meaning domains had notable similarities and differences (**Table 2**). Interestingly the spectro-temporal parameters featured in most of the meaning domains: the FFT in all eight, spectral tilt and the speech envelope in six, suggesting that changes in a pseudoword’s spectral profile over time and the distribution of its power across different frequencies play vital roles underpinning sound-iconic associations. Importantly, the phonetic features related to voicing, manner and place of articulation, and information related to vowel formants are comprehensively represented in the FFT but only partially in spectral tilt. The FFT captures fine-grained energy distribution across the entire frequency range, allowing identification of specific formants and spectral patterns that carry information related to place, manner, and voicing. In contrast, spectral tilt summarizes the overall slope of the spectrum and reflects a more general characterization of the frequency distribution, so it lacks the detailed frequency-specific information over time^83^. The speech envelope captures the alternation between high-energy voiced segments such as sonorants and vowels, with sustained periodic energy, and the transient bursts and amplitude dips of obstruents including stops, fricatives and affricates^84^. In this way, the temporal envelope provides a coarse but informative representation of voicing and manner contrasts, complementing the spectral detail of the FFT and the broader distribution captured by spectral tilt. These features are also significant from a neural standpoint because the initial stages of speech perception involve processing the spectro-temporal properties represented by the FFT and spectral tilt^85,86^. This indicates that spectro-temporal signatures encode rich information that is fundamental for non-arbitrary sound-to-meaning associations spanning multiple meaning domains.

Measures of vocal fold vibration and durational cues represent information fundamental for distinguishing phonological voicing^87^. We observed that the fraction of unvoiced frames, which measures the percentage of time windows that do not engage the vocal folds, featured in the optimal combination of features for six meaning domains, but was absent in the roughness and valence domains (**Table 2**). The fraction of unvoiced frames provides cues about the phonemic content of an utterance^43^, being lower if the utterance includes exclusively voiced elements like sonorants and vowels, and higher if it includes elements like unvoiced obstruents. Such phonemic cues are critical in forming sound-iconic associations^9,19^ and therefore, sound iconicity judgments across multiple domains are sensitive to features reflected in this parameter. However, it is likely that in the roughness and valence domains, the fraction of unvoiced frames was absent because judgments in these domains rely on the richer information captured by the FFT and speech envelope. These features convey detailed spectral and temporal profiles of pseudowords, offering more nuanced cues about the distribution of the component sonorants and obstruents in the pseudowords than the unitary measure provided by the fraction of unvoiced frames.

We found that stimulus duration featured in the optimal model for five domains. Our findings extend and generalize those of Knoeferle et al.^23^ demonstrating the significance of durational cues in sound-size iconicity. Most existing evidence shows how the phonological features that comprise a word determine the nature of its iconic association. Interestingly, the vowels and consonants that represent these phonological features are characterized by distinctive time profiles, with vowel duration being modulated by the consonant environment^88^ and duration influencing the perception of vowel quality^59^. Taken together, the duration of a pseudoword is related to the phonological characteristics comprising the spoken word, and thus provides relevant cues for making sound iconicity judgements. Duration did not feature in the optimal combination for roughness, arousal, and valence, likely because the iconic relationships in these domains depend on parameters that capture the extent of momentary fluctuations and fine-grained spectral and voicing characteristics of sonorants and obstruents, rather than the overall temporal extent or total duration of the sound.

We found that shimmer featured in the optimal combination only for the shape and hardness domains, while the speech envelope was not included in the optimal models for either. This suggests that for the current set of pseudowords, shape and hardness judgments rely more on information related to local amplitude variability, rather than the overall amplitude profile conveyed by the speech envelope.

The HNR features specifically in the optimal combination for the arousal and weight domains. Evidence shows that heightened arousal states, such as stress, reduce HNR through increased vocal fold tension and irregular vibration^89,90^, while higher body weight lowers HNR by altering subglottal pressure and vocal tract damping^91^. These converging effects may explain why HNR features only in the arousal and weight domains, where both emotional and physiological factors directly modulate the harmonic structure of the voice.

The MAC, pulse number, jitter, F0 and F0_SD_ featured in the optimal combination only for the brightness domain. This suggests that these parameters may carry specific information relevant for distinguishing brightness-related sound-iconic judgments, rather than being broadly representative of acoustic-phonetic properties across meaning domains. For instance, both the fraction of unvoiced frames and pulse number capture aspects of phonation or voicing in the pseudoword. However, the fraction of unvoiced frames reflects the relative proportion of unvoiced to voiced periods, whereas pulse number simply reports the number of glottal pulses in the signal. It is possible that for the current set of pseudowords, brightness-related distinctions rely more on absolute measures of voicing (pulse number) than on relative measures (the fraction of unvoiced frames). Similarly, MAC and jitter, which index the regularity of vocal fold vibrations, may be particularly relevant for perceiving fine-grained periodicity or stability cues that listeners associate with brightness, even if these cues are not crucial in other meaning domains. In this way, brightness judgments might rely on a specific set of voicing stability and periodicity features, captured by the MAC, jitter, and pulse number, that do not play the same role for other domains. Whether the brightness-related relevance of these features reflects properties of the pseudoword set, the speaker, or both requires further investigation. The contribution of F0 is consistent with the finding that participants match high-pitched sounds to lighter, and low-pitched sounds to darker, stimuli^70^.

Interestingly, we observed that, except for HNR, all other parameters in the weight model were the same as for the size domain. Notably, Lacey et al.^9^ found that, for the same set of pseudowords as used in the present study, heaviness ratings were correlated with bigness ratings, and lightness with smallness. The shared acoustic parameters might stem from the fact that weight and size mostly co-vary, i.e. big objects in most cases are heavier than small objects. However, it is important to note that this relationship is not absolute, since weight also depends on density. As a result, a large but low-density object can weigh less than a smaller, high-density one.

### Sound iconicity ratings of real words

Extending our KNN approach, we demonstrated that the predicted shape ratings for real words, derived from pseudowords in their acoustic neighborhoods, were strongly correlated with their perceptual ratings. Although linguistic corpus studies demonstrate instances of *iconicity* in natural languages, where a word’s form somehow bears resemblance to its meaning, the vast majority of these studies only examine whether iconicity exists in the first place^32,92^. The ratings obtained this way are not divided into specified meaning domains and more importantly, these ratings are prone to influence by subjective semantic knowledge. Conversely, our approach allows us to use pseudowords to first narrow down the most relevant acoustic parameters that aid in sound-iconic associations in a given meaning domain, and then, based on those acoustic parameters, to find the nearest pseudoword neighbors for each real word. The real word rating is determined by the mode of the perceptual ratings of these nearest pseudoword neighbors. Following this procedure allowed us to apply the principles emerging from pseudowords to generate ratings for real words, free of any semantic bias. These predicted ratings were then compared to perceptual shape ratings for the real words; the two sets of ratings were significantly correlated. The perceptual ratings of the real words may, to some extent, reflect listeners’ semantic expectations, since they already know the meanings of these words, rather than sensitivity to acoustic features alone. Thus, we cannot currently rule out the possibility that observed correspondence between ratings of pseudowords and real words may partially be due to semantic influences: our approach substantially reduces the potential for semantic bias, but it cannot completely eliminate it.

### Translating iconicity into practice

Our KNN-based approach represents a principled way to empirically identify the acoustic parameters underlying sound-iconic judgements. The approach depends on the premise that, in certain parameter spaces, data points belonging to similar categories are closer to each other than data points that belong to different categories, analogous to our natural ability to form categories and classify items based on specific sets of features^93,94^. In addition, the operational principle of the KNN algorithm makes our approach transparent to the underlying relationships learned by the acoustic parameter models, as opposed to the general opacity of widely employed machine-learning approaches like SVMs and deep learning methods.

Above all, the most significant contribution of our approach is that it offers a way to employ the principles of iconic associations, deduced from pseudowords, to measure the degree of sound iconicity in natural languages. Notably, since the measures are based on the acoustic features that drive sound-iconicity associations in pseudowords, they are not influenced by the conventional meaning of the words in that language. This has potential implications in the study of language development and applications in the field of consumer psychology.

Research suggests that during the initial stages of acquiring a vocabulary, iconicity may provide an effective, even essential, scaffold that facilitates early acquisition of words in that vocabulary^79,95^. Interestingly, in Japanese, iconic words, especially those which refer to action, are used more in speech by and toward children, and children are sensitive to the meanings of sound-iconic verbs^96^. In light of such evidence, our findings offer avenues for studying the impact of different acoustic features, fundamental for iconic associations in a particular domain, that affect the learning of vocabulary of that domain. In addition, given the significance of iconicity in early language development^96,97^, a potential remedy for children facing difficulty in language learning could involve adjusting the inputs given to them, to include more iconic words. Although recent work suggests that the role of iconicity in early stages of language acquisition may be more limited than previously thought^98^, we propose, nonetheless, that sound-iconic cues may still provide a valuable scaffold for preschool children experiencing language-learning difficulties, whether or not they play a primary role in typically developing infants. To this end, our approach could potentially aid in finding an iconically rich vocabulary that could guide parents and caregivers of children facing difficulty in language learning. Similar considerations apply to rehabilitation of individuals with acquired language difficulties, e.g. in post-stroke aphasia, where sound-iconic associations seem to be preserved^99,100^.

Research in consumer psychology shows that iconic brand names can convey product attributes like size or strength, shaping favorable impressions even without explicit marketing^101,102^. While prior work focused on broad contrasts (e.g., rounded vs unrounded vowels)^103^, our findings offer a roadmap for finer control, suggesting that rounder or larger products are best expressed with longer names emphasizing higher power in lower frequencies.

## Conclusions

In sum, our findings indicate that different acoustic parameters underlie the iconic associations of pseudoword sounds in different domains. In addition, we present an approach that can be used to assess the iconicity of spoken words in natural languages, unconfounded by semantic bias. Further research can extend our findings and methods to understand the roles of these acoustic parameters in natural language learning and processing. In addition, future neuroimaging and electrophysiological studies can provide insight into how the patterns represented in these acoustic parameters are represented neurally, which could pave the way to understanding the neural basis of sound iconicity.

## ACKNOWLEDGEMENTS

This work was supported by grants to KS and LCN from the National Eye Institute at the NIH (R01EY025978) and the Emory University Research Council, and institutional funds provided to KS by Penn State College of Medicine. Earlier versions of these data were presented at the 2022 meeting of the Cognitive Neuroscience Society (Hoffmann et al., 2022; Nygaard et al., 2022). A previous version of this manuscript was posted as a preprint on bioRxiv (https://doi.org/10.1101/2024.09.27.615393). We thank Saachi Nayak for data collection on the roughness aspect of texture from her MS thesis under the direction of KS and SL (Nayak, 2024).

## AUTHOR CONTRIBUTIONS

SL, KS, LCN and GVK designed research; JD collected data; GVK analyzed data; GVK, SL, KS and LCN wrote the paper.

## DATA AVAILABILITY

Stimuli, rating data and scripts used for analyses in this study are available at https://osf.io/hdm7w/

