## Supplementary Material for "Acoustic parameter combinations underlying mapping of pseudoword sounds to multiple domains of meaning: representational similarity analyses and machine-learning models"

G. Vinodh Kumar[1], Simon Lacey[1,2,3], Josh Dorsi[1], Lynne C. Nygaard[4,*] & K. Sathian[1,2,3,*]

Departments of Neurology[1], Neuroscience & Experimental Therapeutics[2],
Penn State College of Medicine, Hershey, PA 17033, USA.
Department of Psychology [3], Penn State College of Liberal Arts,
University Park, PA 16802, USA
Department of Psychology[4], Emory University, Atlanta, GA 30322, USA

Corresponding authors*:

K. Sathian
Department of Neurology
Penn State Health Milton S. Hershey Medical Center
Hershey, PA 17033-0859, USA


Lynne C. Nygaard
Department of Psychology
Emory University
College of Arts and Sciences
Atlanta, GA 30322, USA


**Supplementary Methods**

### I. Computation of speech envelope, spectral tilt, and FFT

The speech envelope was calculated using the root mean square of the amplitude and a sliding window length of 120 data points; the window was centered over each data point and moved one data point at a time along the vector of 10077 data points, thus giving 10077 measurements per pseudoword. The pairwise dissimilarities between pseudowords were computed using Pearson correlations.

The calculation of any spectral parameter is limited by the Nyquist frequency (0.5 x sampling rate) to avoid aliasing, or false signals[1]. To calculate spectral tilt, we first generated the power spectral density function over the 10077 data points of the normalized pseudoword duration. The resulting power spectral density is mirror-symmetrical along the full vector of 10077 datapoints: applying the Nyquist frequency limit reduces this to a vector with length 5038 and removes the symmetry. Over the vector of 5038 datapoints, we used a sliding window of 600 data points with overlap of 15, i.e, after the first 600 data points, the window moves forward 585 data points until it goes past the end of the vector. The final, incomplete window was discarded. Within each window, we fitted a polynomial to the 600 data points and estimated the slope from the polynomial coefficient of the periodogram (power across frequency). This provided 8 measurement windows, corresponding to 8 slopes, providing a measure of spectral tilt over time. Note that at the end of the vector, the final, incomplete window was 343 data points; discarding this represents a loss of 6.8% of the data. Since there were only 8 measurements for each pseudoword, the pairwise dissimilarities between pseudowords were computed using Spearman correlations.

The FFT was calculated with a sliding time window of 100 data points with an overlap of 80, and a frequency window of 100 discrete Fourier transform points from 1 to 11 kHz. The output of the spectrogram function in MATLAB was a vector of length 499 for each frequency; since we only used a limited range of the frequencies, i.e., 1-11 kHz, the vectors were concatenated, resulting in 5489 measurements per pseudoword. The pairwise dissimilarities between pseudowords were computed using Pearson correlations.

### II. Sensitivity analysis of k across meaning domains

The KNN algorithm described in Section 2.6 of the main manuscript was repeated with the same procedure, except that the neighborhood size k was systematically varied. Specifically, k was set to approximately 5% (26), 7.5% (40), and 10% (53) of the total number of pseudowords (537) to assess whether the same parameter combination identified for k = 23 would also emerge across different k values. This analysis was conducted for two meaning domains: size and arousal, because they yielded the lowest and highest correlations, respectively, between the perceptual ratings and the ratings generated by the algorithm.

We found that that the optimal feature combinations remained largely consistent across different k values, particularly with respect to the spectro-temporal parameters and the voice parameter fraction of unvoiced frames – these parameters occurred most consistently in the models across domains (**Supplementary Table 1**).

### III. Stability analysis of parameter combinations

To assess the stability of the optimal parameter combinations with respect to variability in the participant sample, we conducted a subsampling analysis within each meaning domain. For this, the KNN algorithm was rerun multiple times while progressively increasing the proportion of participants included in the training set in increments of 10%. Specifically, random subsets of participants were drawn to represent 10%, 20%, 30%, …, 90% of the total participant pool. For each sampling proportion, 100 independent random draws were performed. The KNN analysis was then applied to each subset, and the optimal parameter combination (as defined in Section 2.6 in the main text) was identified for each run. This allowed us to asses assess how robust the parameter selection was to reductions in the number of participants.

We observed that the models showed consistent performance with different subsamples (**Supplementary Figures 1-8**).

**Supplementary Table 1:** Optimal parameter combinations across varying number of neighbors (k = 5%, 7.5%, 10% of 537 pseudowords) for the size and arousal domains. The table shows the best-performing combination for each k value, allowing comparison with the original k = 23 results.

FFT: Fast Fourier transform, ST: Spectral tilt, SE: Speech envelope, PN: Pulse number, FUF: Fraction of unvoiced frames, HNR: Harmonics-to-noise ratio, MAC: Mean autocorrelation

| | | **Optimal combinations** | |
|---|---|---|---|
| %age | k | ***Size*** | ***Arousal*** |
| | **23** | FFT, ST, FUF, shimmer and duration | FFT, ST, SE, FUF and HNR |
| 5% | **26** | FFT, ST, SE, FUF and duration | FFT, ST, SE, FUF, MAC and duration |
| 7.5% | **40** | FFT, ST, SE, PN, FUF, shimmer, $F0_{SD}$ and F0. | FFT, ST, SE, FUF, PN, HNR and duration |
| 10% | **53** | FFT, ST, FUF, HNR, $F0_{SD}$ and duration | FFT, ST, SE, FUF, HNR, F0 and duration |

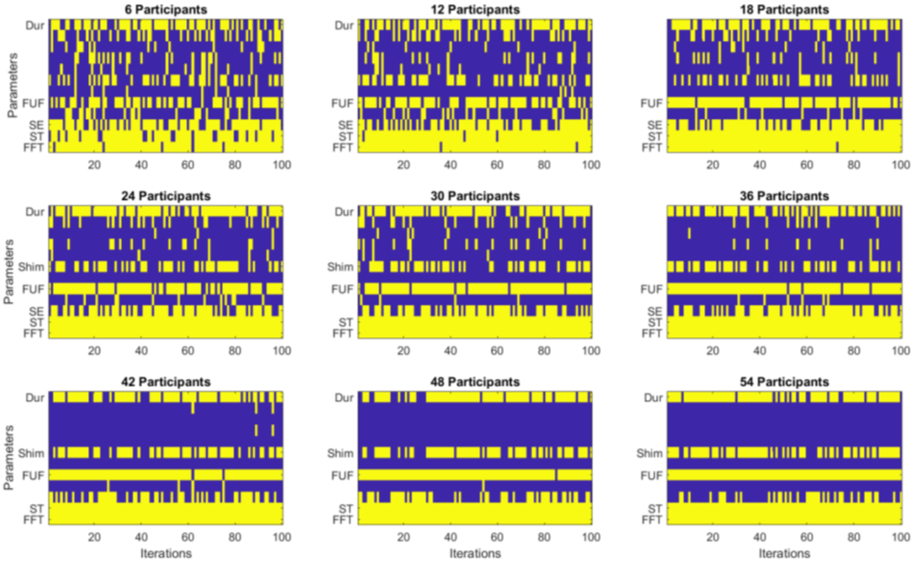


**Supplementary Figure 1: Stability of optimal parameter combination for Shape across participant subsampling proportions.** Each subplot shows the results of a subsampling analysis in which the KNN algorithm was rerun 100 times on random subsets of participants, with the subset size increasing from 10% to 90% of the total sample in increments of 10% (rows from top left to bottom right). Within each subplot, columns represent independent iterations and rows represent individual acoustic parameters. Yellow cells indicate the presence of a parameter in the optimal combination for that iteration, whereas blue cells indicate its absence. The parameters listed on each subplot are the most frequent combination for that particular subset of participants. This visualization illustrates how consistently each parameter appears in the optimal set as the number of participants varies, and also provides the consistency of optimal combination of parameters, providing an index of robustness to sample size variation.

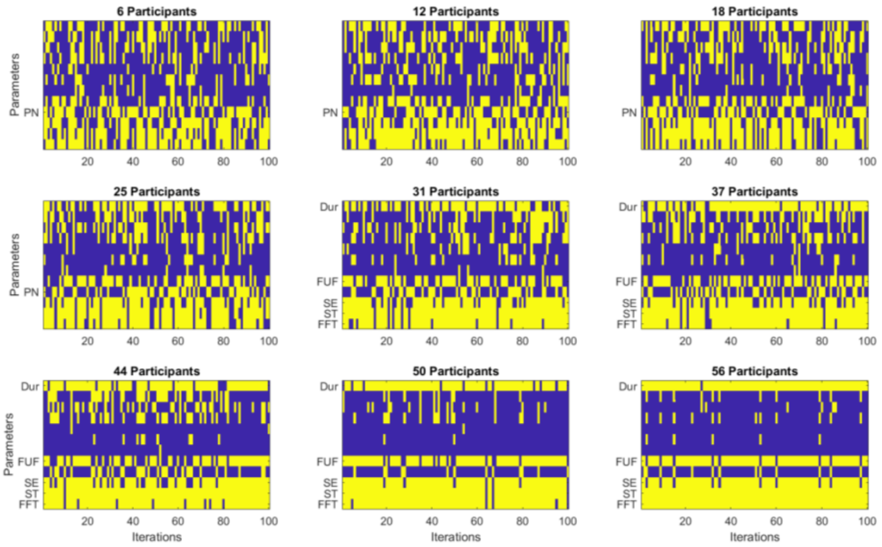


**Supplementary Figure 2: Stability of optimal parameter combination for Size across participant subsampling proportions.** Each subplot shows the results of a subsampling analysis in which the KNN algorithm was rerun 100 times on random subsets of participants, with the subset size increasing from 10% to 90% of the total sample in increments of 10% (rows from top left to bottom right). All other details as for Supplementary Fig. 1

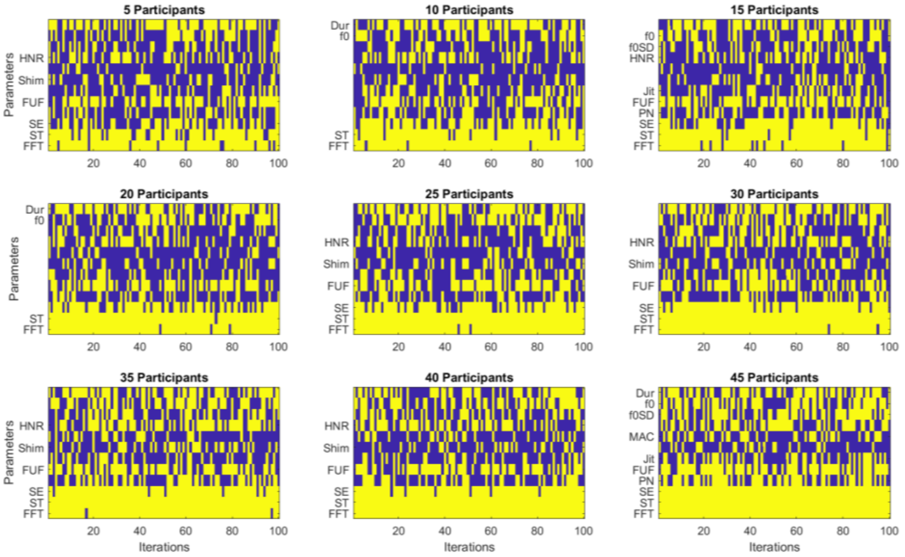


**Supplementary Figure 3: Stability of optimal parameter combination for Brightness across participant subsampling proportions.** Each subplot shows the results of a subsampling analysis in which the KNN algorithm was rerun 100 times on random subsets of participants, with the subset size increasing from 10% to 90% of the total sample in increments of 10% (rows from top left to bottom right). All other details as for Supplementary Fig. 1

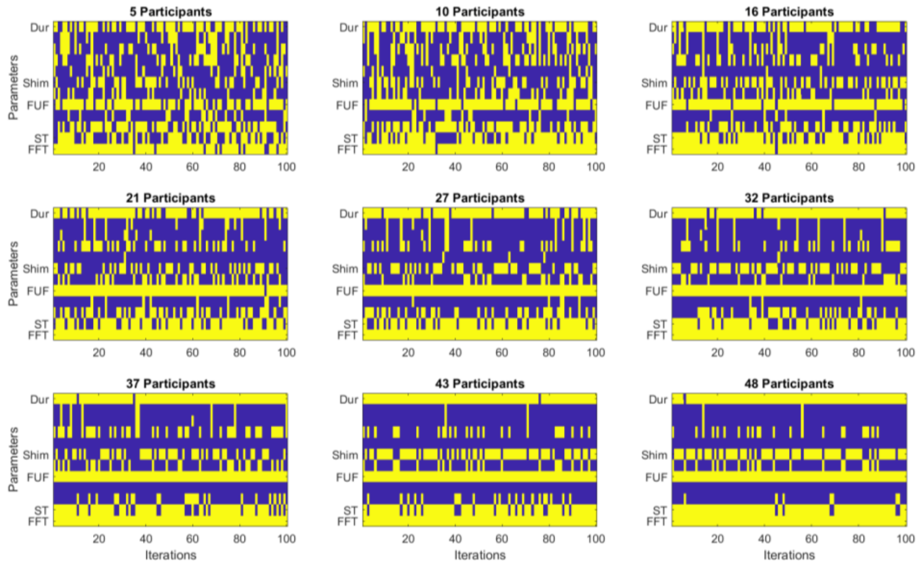


**Supplementary Figure 4: Stability of optimal parameter combination for Hardness across participant subsampling proportions.** Each subplot shows the results of a subsampling analysis in which the KNN algorithm was rerun 100 times on random subsets of participants, with the subset size increasing from 10% to 90% of the total sample in increments of 10% (rows from top left to bottom right). All other details as for Supplementary Fig. 1

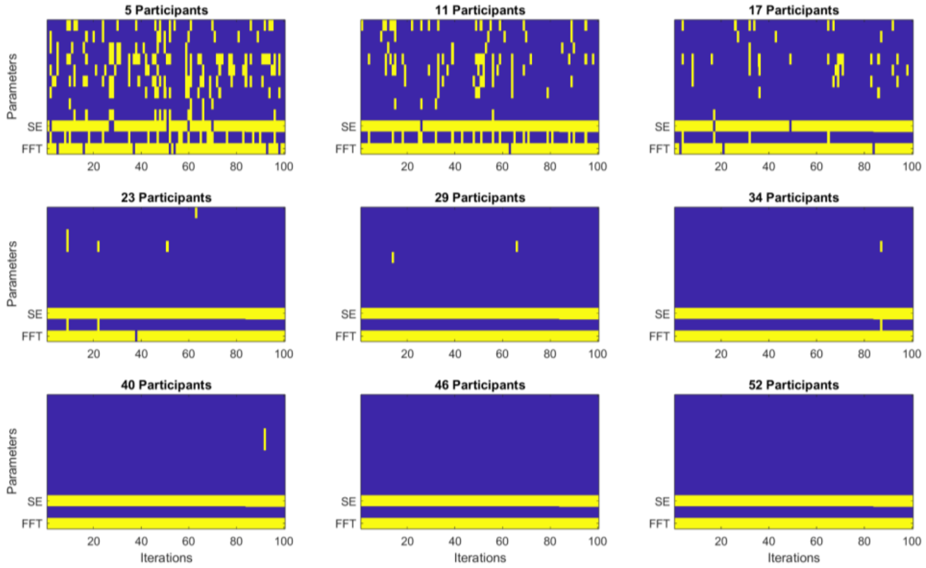


**Supplementary Figure 5: Stability of optimal parameter combination for Roughness across participant subsampling proportions.** Each subplot shows the results of a subsampling analysis in which the KNN algorithm was rerun 100 times on random subsets of participants, with the subset size increasing from 10% to 90% of the total sample in increments of 10% (rows from top left to bottom right). All other details as for Supplementary Fig. 1

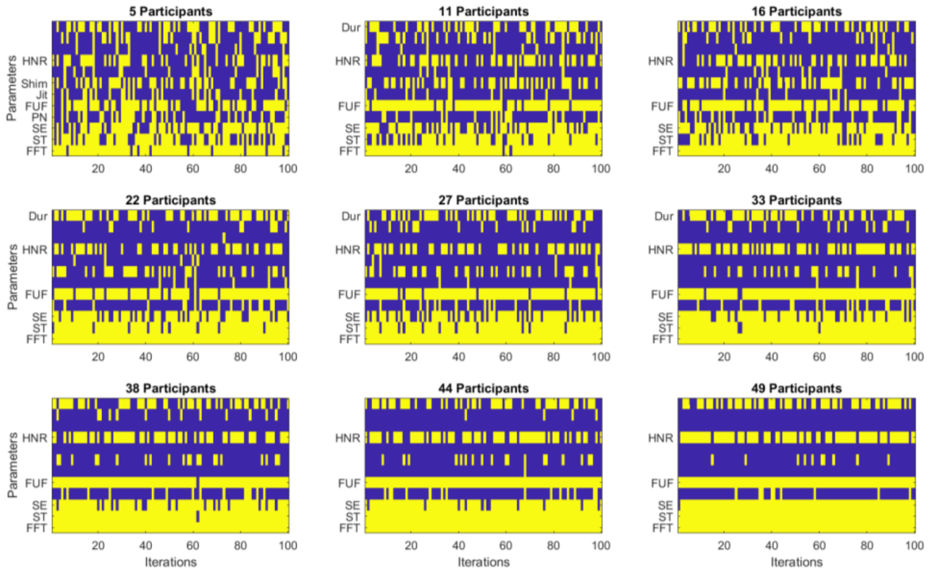


**Supplementary Figure 6: Stability of optimal parameter combination for Weight across participant subsampling proportions.** Each subplot shows the results of a subsampling analysis in which the KNN algorithm was rerun 100 times on random subsets of participants, with the subset size increasing from 10% to 90% of the total sample in increments of 10% (rows from top left to bottom right). All other details as for Supplementary Fig. 1

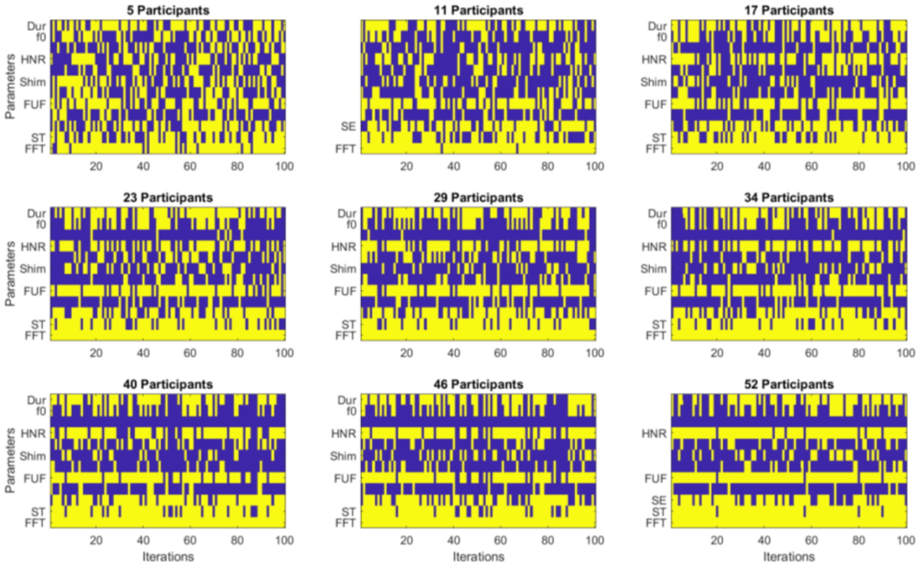


**Supplementary Figure 7: Stability of optimal parameter combination for Arousal across participant subsampling proportions.** Each subplot shows the results of a subsampling analysis in which the KNN algorithm was rerun 100 times on random subsets of participants, with the subset size increasing from 10% to 90% of the total sample in increments of 10% (rows from top left to bottom right). All other details as for Supplementary Fig. 1

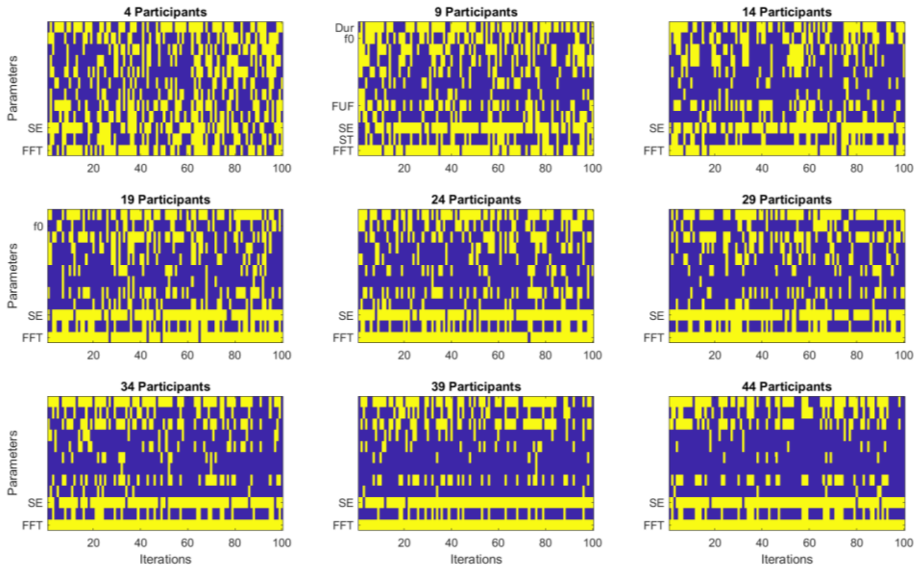


**Supplementary Figure 8: Stability of optimal parameter combination for Valence across participant subsampling proportions.** Each subplot shows the results of a subsampling analysis in which the KNN algorithm was rerun 100 times on random subsets of participants, with the subset size increasing from 10% to 90% of the total sample in increments of 10% (rows from top left to bottom right). All other details as for Supplementary Fig. 1

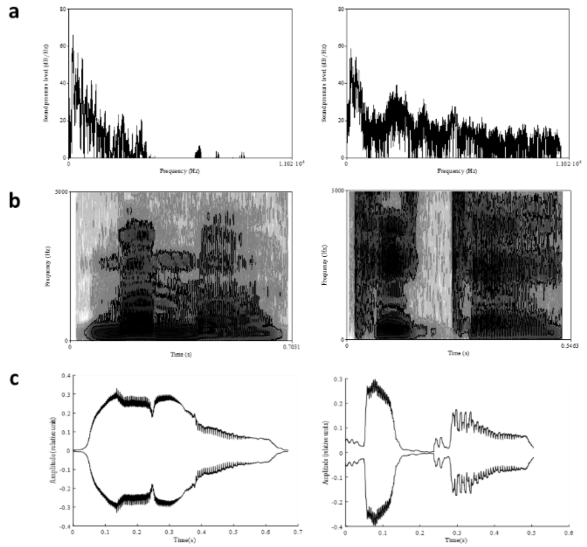


**Supplementary Figure 9:** Spectro-temporal properties of representative highly-rated rounded (/mʊmo/, mean rating 2.72, left column) and pointed (/tike/, mean rating 5.5, right column) pseudowords, ordered by the strength of their relationship to pseudoword ratings, strongest to weakest: (a) spectral tilt: frequency low to high, left to right; (b) spectrogram: frequency low to high, bottom to top; shading = power, low power lighter and high power darker; (c) speech envelope.

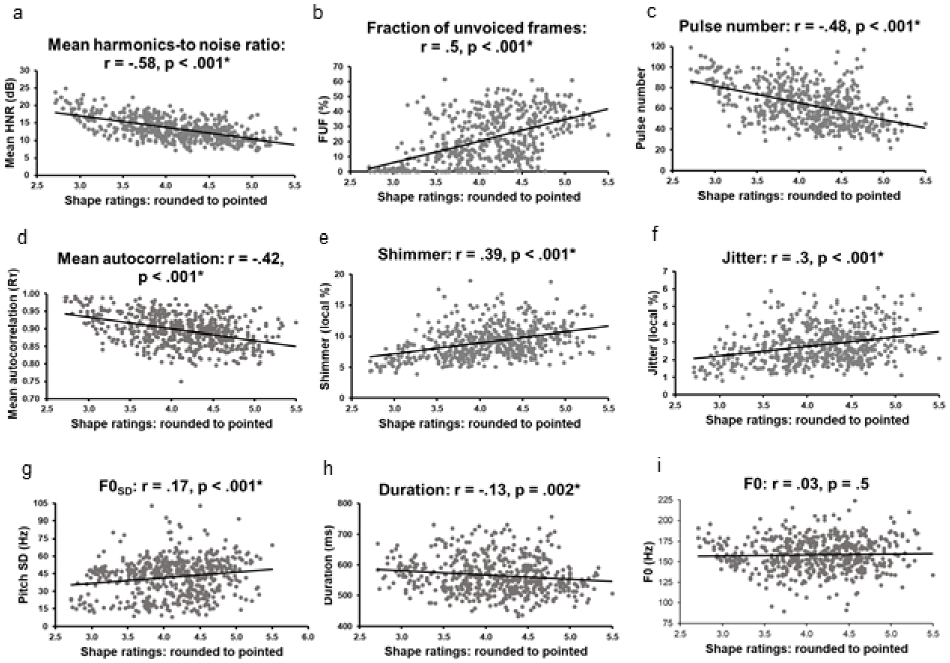


**Supplementary Figure 10:** Correlations between auditory perceptual ratings for shape and nine voice-quality parameters, ordered, top to bottom, left to right, from the strongest to the weakest relationship, whether positive or negative. r = Pearson correlation coefficient; *correlation passes the Bonferroni-corrected α of 0.0056 for nine tests; df = 535 for all nine voice parameters.

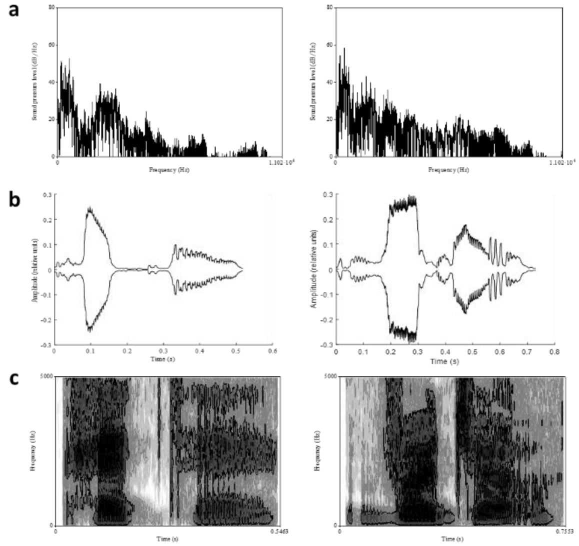


**Supplementary Figure 11:** Spectro-temporal properties of representative highly-rated small (/pete/, mean rating 3.41, left column) and big (/vʊdʒo/, mean rating 5.06 right column) pseudowords, ordered by the strength of their relationship to pseudoword ratings, strongest to weakest, top to bottom. (a) spectral tilt; (b) speech envelope; (c) spectrogram. Axis labels etc., as for Supplementary Figure 9

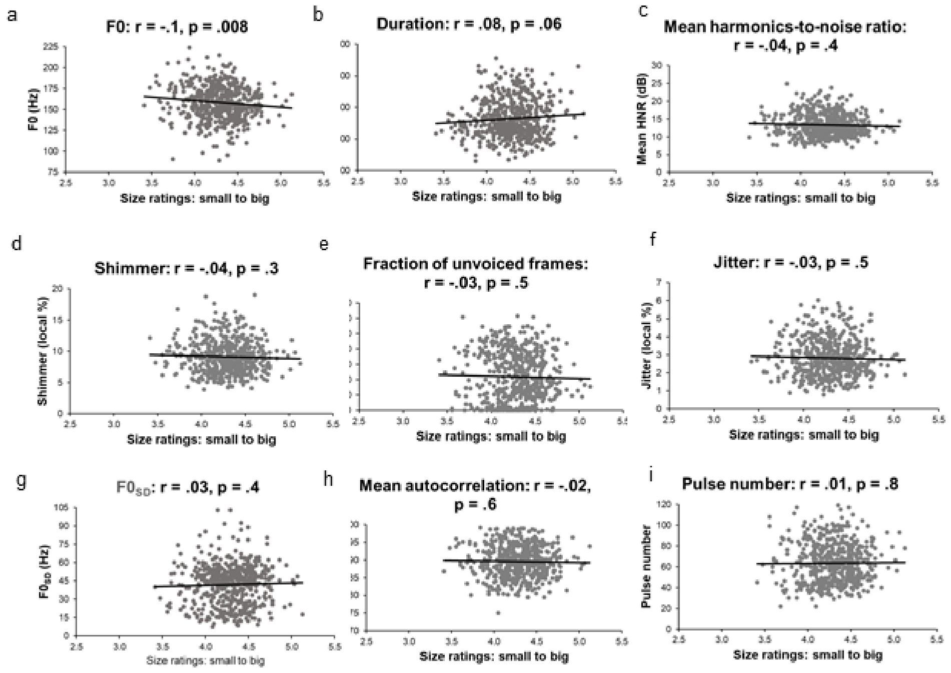


**Supplementary Figure 12:** Correlations between auditory perceptual ratings for size and nine voice-quality parameters, ordered, top to bottom, left to right, from the strongest to the weakest relationship, whether positive or negative. r, *, and df as for Supplementary Figure 10.

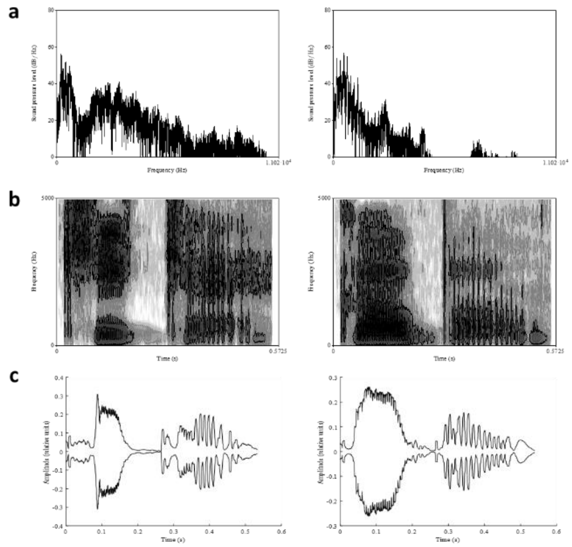


**Supplementary Figure 13:** Spectro-temporal properties of representative highly-rated bright (/kike/, mean rating 2.86, left column) and dark (/gobo/, mean rating 5.08, right column) pseudowords, ordered by the strength of their relationship to pseudoword ratings, strongest to weakest, top to bottom. (a) spectral tilt; (b) spectrogram; (c) speech envelope. Axis labels etc. as for Supplementary Figure 9.

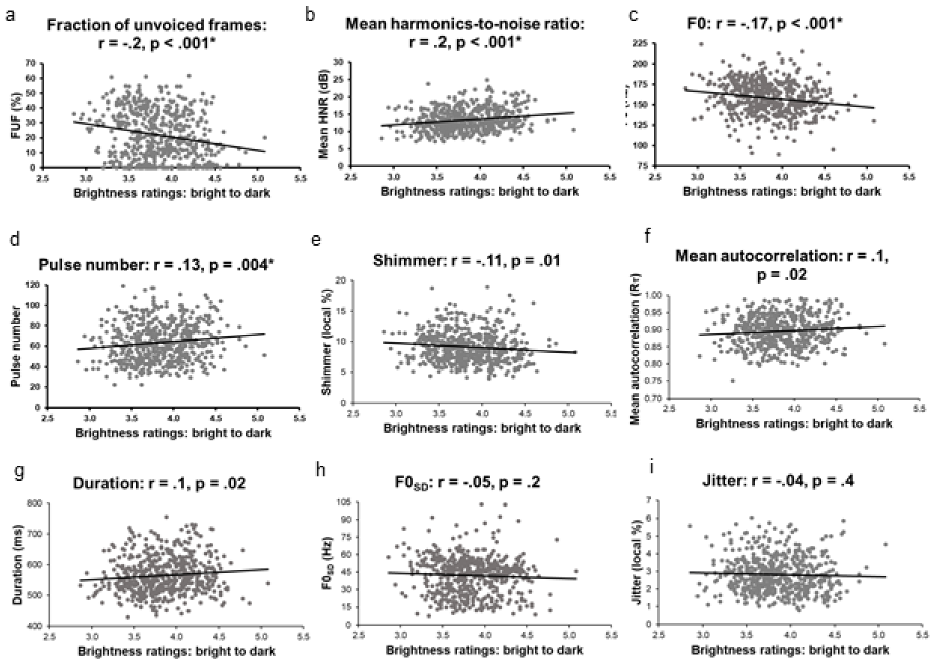


**Supplementary Figure 14:** Correlations between auditory perceptual ratings for brightness and nine voice-quality parameters, ordered, top to bottom, left to right, from the strongest to the weakest relationship, whether positive or negative. r, *, and df as for Supplementary Figure 10.

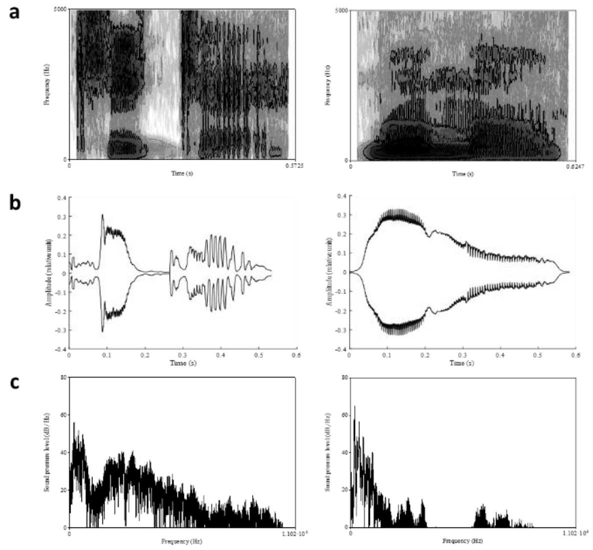


**Supplementary Figure 15:** Spectro-temporal properties of representative highly-rated hard (/kike/, mean rating 2.85, left column) and soft (/mumo/, mean rating 5.07, right column) pseudowords, ordered by the strength of their relationship to pseudoword ratings, strongest to weakest, top to bottom. (a) spectrogram; (b) speech envelope; (c) spectral tilt. Axis labels etc., as for Supplementary Figure 9.

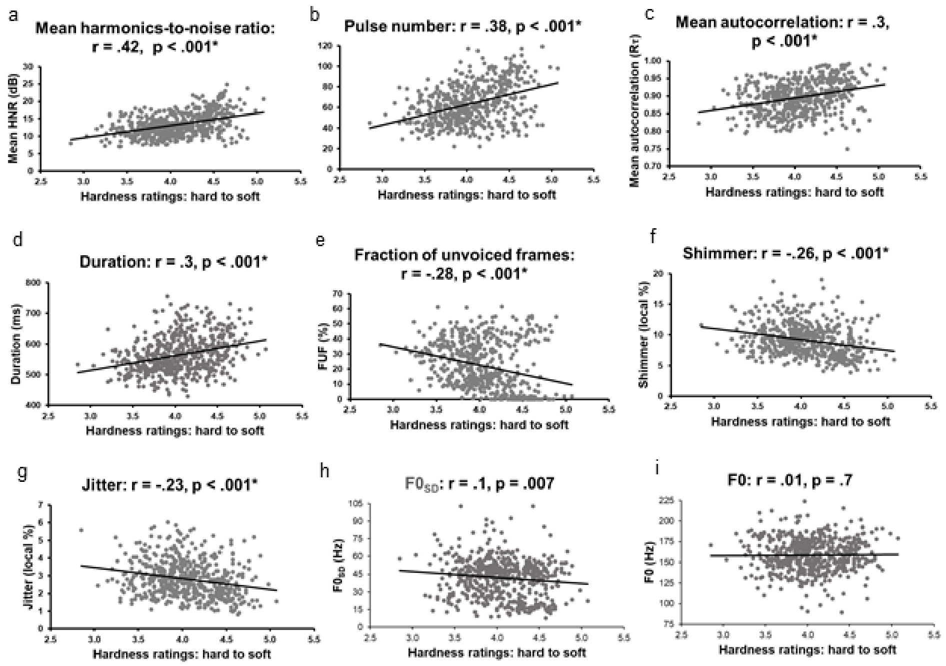


**Supplementary Figure 16:** Correlations between auditory perceptual ratings for hardness and nine voice-quality parameters, ordered, top to bottom, left to right, from the strongest to the weakest relationship, whether positive or negative. r, *, and df as for Supplementary Figure 10.

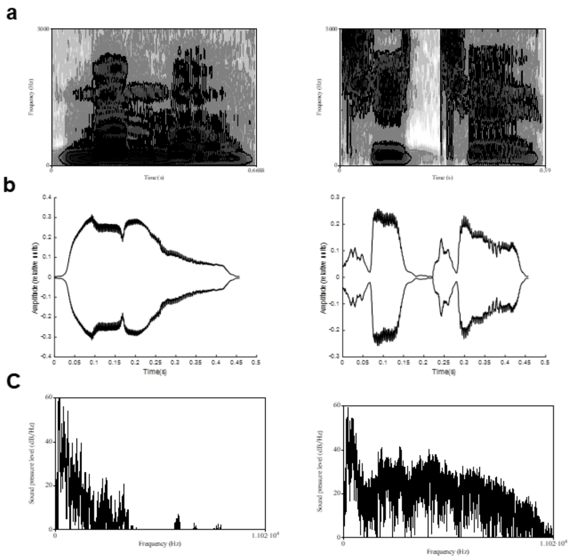


**Supplementary Figure 17:** Spectro-temporal properties of representative highly rated smooth (/mumo/, mean rating 2.33, left column) and highly-rated rough (/tʃitʃe/, mean rating 4.09, right column) pseudowords, ordered by the strength of their relationship to pseudoword ratings, strongest to weakest, top to bottom. (a) spectrogram; (b) speech envelope; (c) spectral tilt. Axis labels etc., as for Supplementary Figure 9.

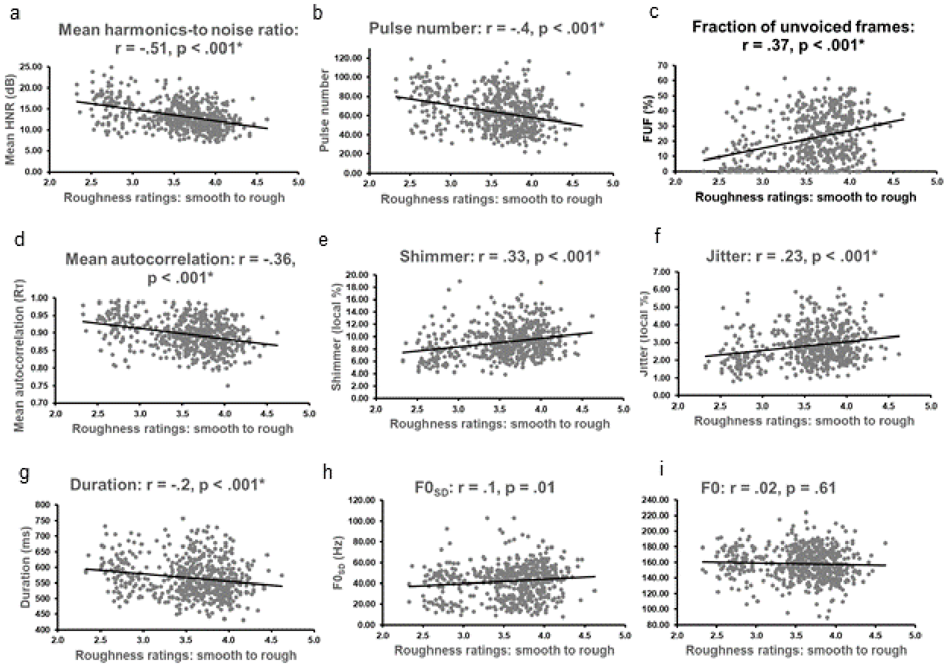


**Supplementary Figure 18:** Correlations between auditory perceptual ratings for roughness and nine voice-quality parameters, ordered, top to bottom, left to right, from the strongest to the weakest relationship, whether positive or negative. r, *, and df as for Supplementary Figure 10.

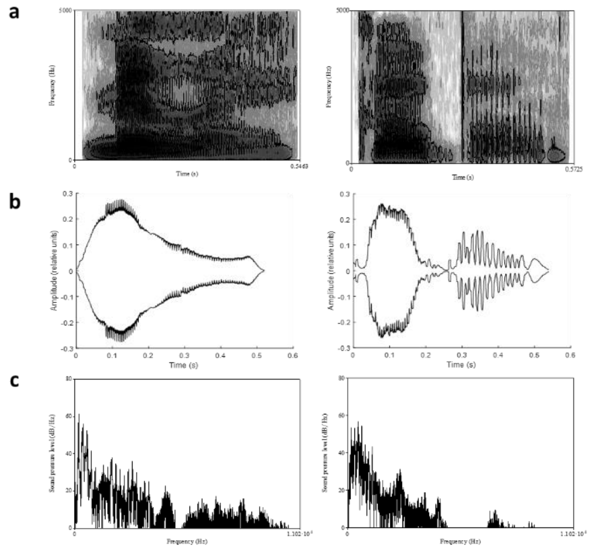


**Supplementary Figure 19:** Spectro-temporal properties of representative highly-rated light (/mIle/, mean rating 3.29, left column) and heavy (/gobo/, mean rating 4.73, right column) pseudowords, ordered by the strength of their relationship to pseudoword ratings, strongest to weakest, top to bottom. (a) spectrogram; (b) speech envelope; (c) spectral tilt. Axis labels etc., as for Supplementary Figure 9.

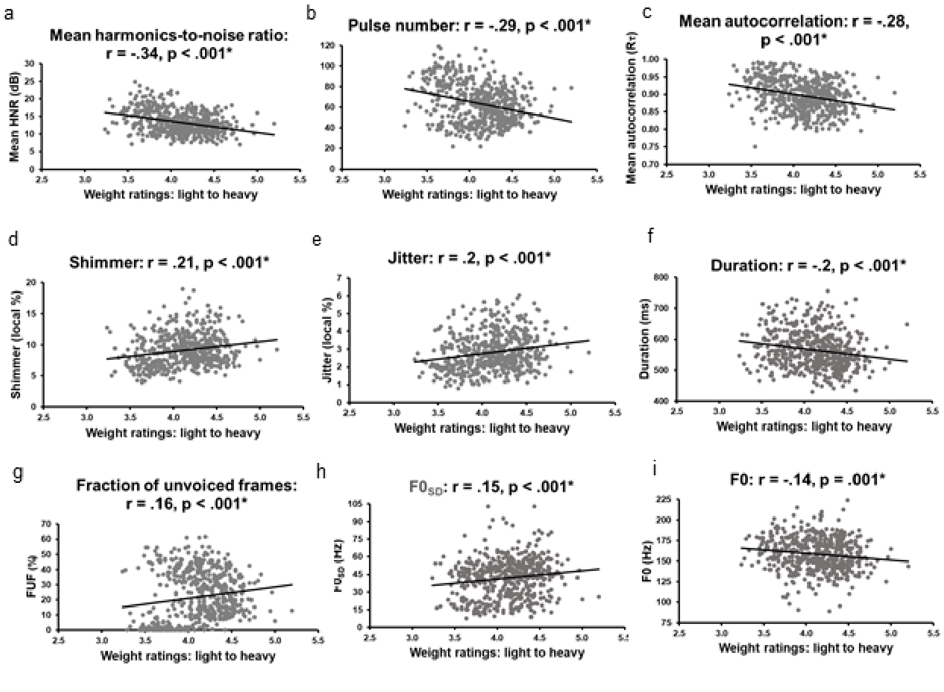


**Supplementary Figure 20:** Correlations between auditory perceptual ratings for weight and nine voice-quality parameters, ordered, top to bottom, left to right, from the strongest to the weakest relationship, whether positive or negative. r, *, and df as for Supplementary Figure 10.

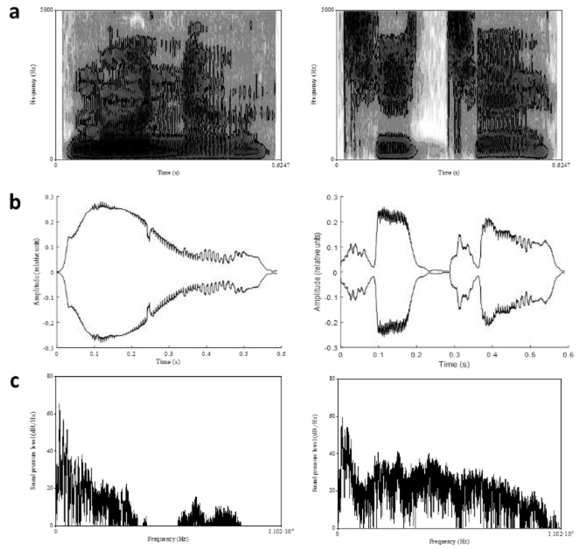

**Supplementary Figure 21:** Spectro-temporal properties of representative highly-rated calming (/munu/, mean rating 2.95, left column) and exciting (/tʃitʃe/, mean rating 4.45, right column) pseudowords, ordered by the strength of their relationship to pseudoword ratings, strongest to weakest, top to bottom. (a) spectrogram; (b) speech envelope; (c) spectral tilt. Axis labels etc., as for Supplementary Figure 9.

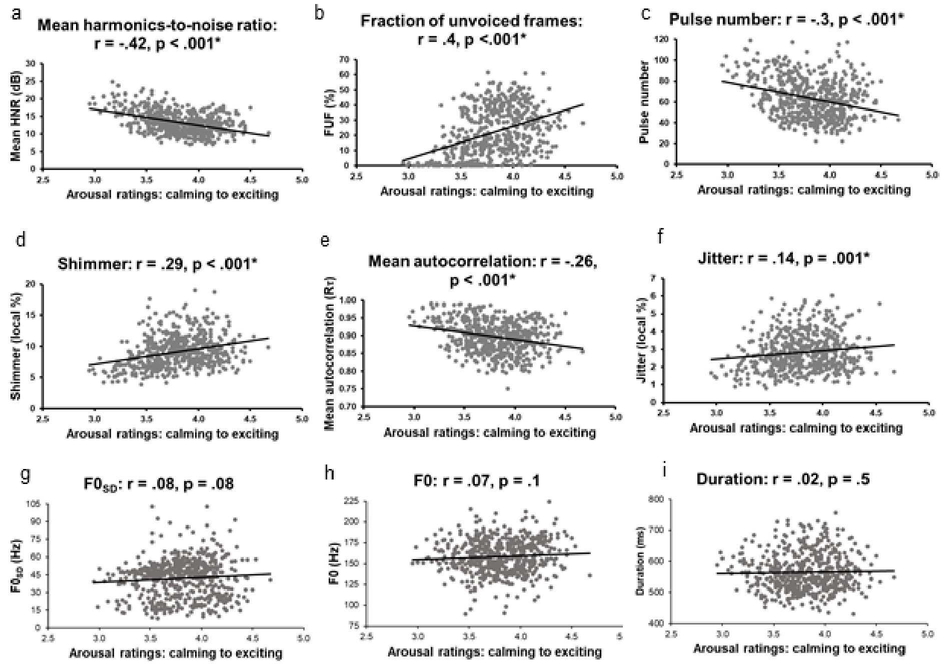


**Supplementary Figure 22:** Correlations between auditory perceptual ratings for arousal and nine voice-quality parameters, ordered, top to bottom, left to right, from the strongest to the weakest relationship, whether positive or negative. r, *, and df as for Supplementary Figure 10.

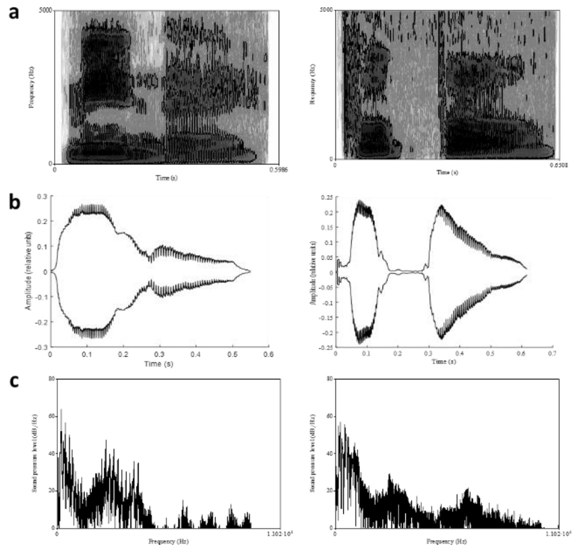


**Supplementary Figure 23:** Spectro-temporal properties of representative highly-rated good (/mine/, mean rating 2.65, left column) and bad (/pupo/, mean rating 4.51, right column) pseudowords, ordered by the strength of their relationship to pseudoword ratings, strongest to weakest, top to bottom. (a) spectrogram; (b) speech envelope; (c) spectral tilt. Axis labels etc., as for Supplementary Figure 9.

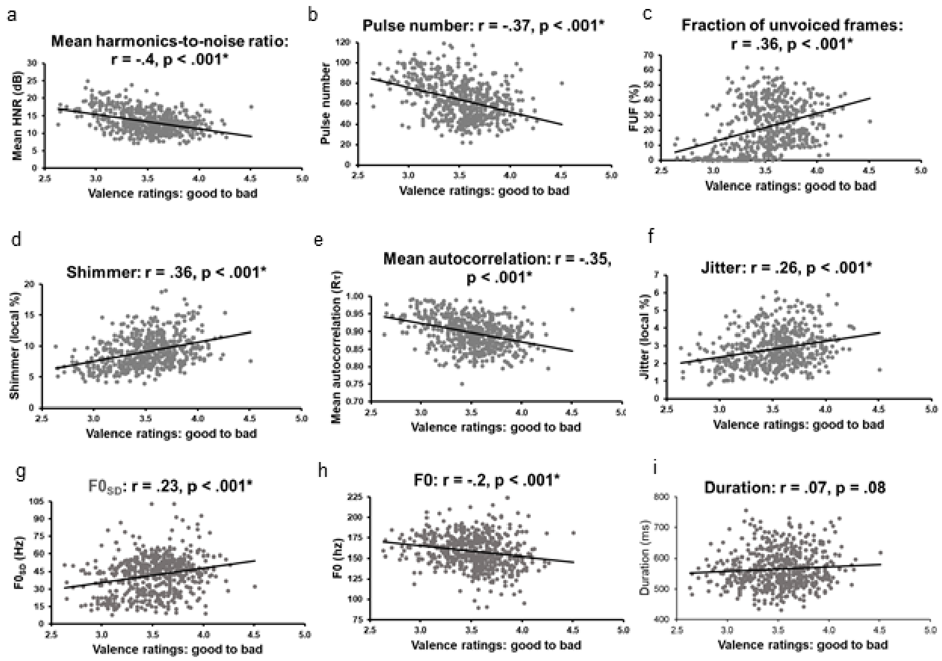


**Supplementary Figure 24:** Correlations between auditory perceptual ratings for valence and nine voice-quality parameters, ordered, top to bottom, left to right, from the strongest to the weakest relationship, whether positive or negative. r, *, and df as for Supplementary Figure 10.

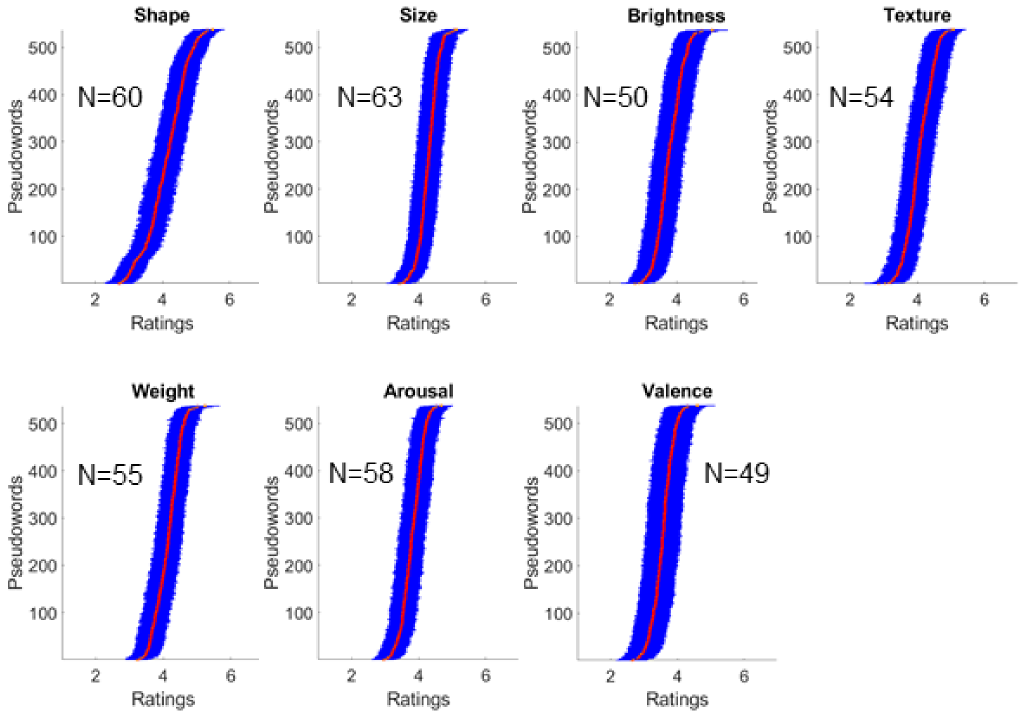


**Supplementary Figure 25:** Variability of perceptual ratings across seven meaning domains for pseudowords. Red dots indicate the mean rating for each pseudoword, while blue vertical lines represent the range corresponding to 95% confidence interval. The order of pseudowords differs across domains and is sorted as follows: Shape: rounded to pointed; Size: small to big; Brightness: bright to dark; Texture: hard to soft; Weight: light to heavy; Arousal: calming to exciting; Valence: good to bad. The number of participants (N) for each domain is indicated in the corresponding subplot.
